# Bacteria use spatial sensing to direct chemotaxis on surfaces

**DOI:** 10.1101/2024.02.13.580113

**Authors:** James H. R. Wheeler, Kevin R. Foster, William M. Durham

## Abstract

Planktonic bacteria navigate chemical gradients using temporal sensing to detect changes in concentration over time as they swim. Here we show that surface-attached bacteria use a fundamentally different mode of sensing during chemotaxis. We combined microfluidic experiments, massively parallel cell tracking, and fluorescent reporters to study how *Pseudomonas aeruginosa* senses chemical gradients during pili-based “twitching” chemotaxis on surfaces. First, we asked whether surface-attached cells use temporal sensing by exposing them to temporal chemical gradients generated via Taylor-Aris dispersion. However, we find that temporal changes in concentration do not induce changes in motility, indicating that twitching cells do not sense chemical gradients like swimming bacteria do. We, therefore, designed experiments to test whether cells can detect chemical gradients across the length of their bodies. In these experiments, we follow the localisation of a fluorescent protein fusion to quantify the chemotactic behaviour of stationary cells in an alternating chemical gradient. We find that *P. aeruginosa* cells can directly sense differences in concentration across the lengths of their bodies, even in the presence of strong temporal fluctuations. Our work reveals that *P. aeruginosa* cells are capable of spatial sensing, thus overturning the widely held notion that bacterial cells are too small to directly sense chemical gradients in space.

## Introduction

Cellular chemotaxis, the ability to sense chemical gradients and actively direct motility along them, plays a central role in many important processes including disease [1, 2], foraging [3, 4], sexual reproduction [5] and multicellular development [6, 7]. There are two distinct ways that cells can sense chemical gradients (**Fig. 1**). Cells using temporal sensing measure changes in chemical concentration over time as they travel along gradients. In contrast, cells using spatial sensing directly compare the concentration of a chemical at different positions along their cell body, independently from cell movement. The two sensing mechanisms are not necessarily mutually exclusive; in some complex signal transduction systems (for example, in certain eukaryotic cells that travel along surfaces using ameboid movement), they can also be used in combination to guide chemotaxis [8].

**Figure 1.**
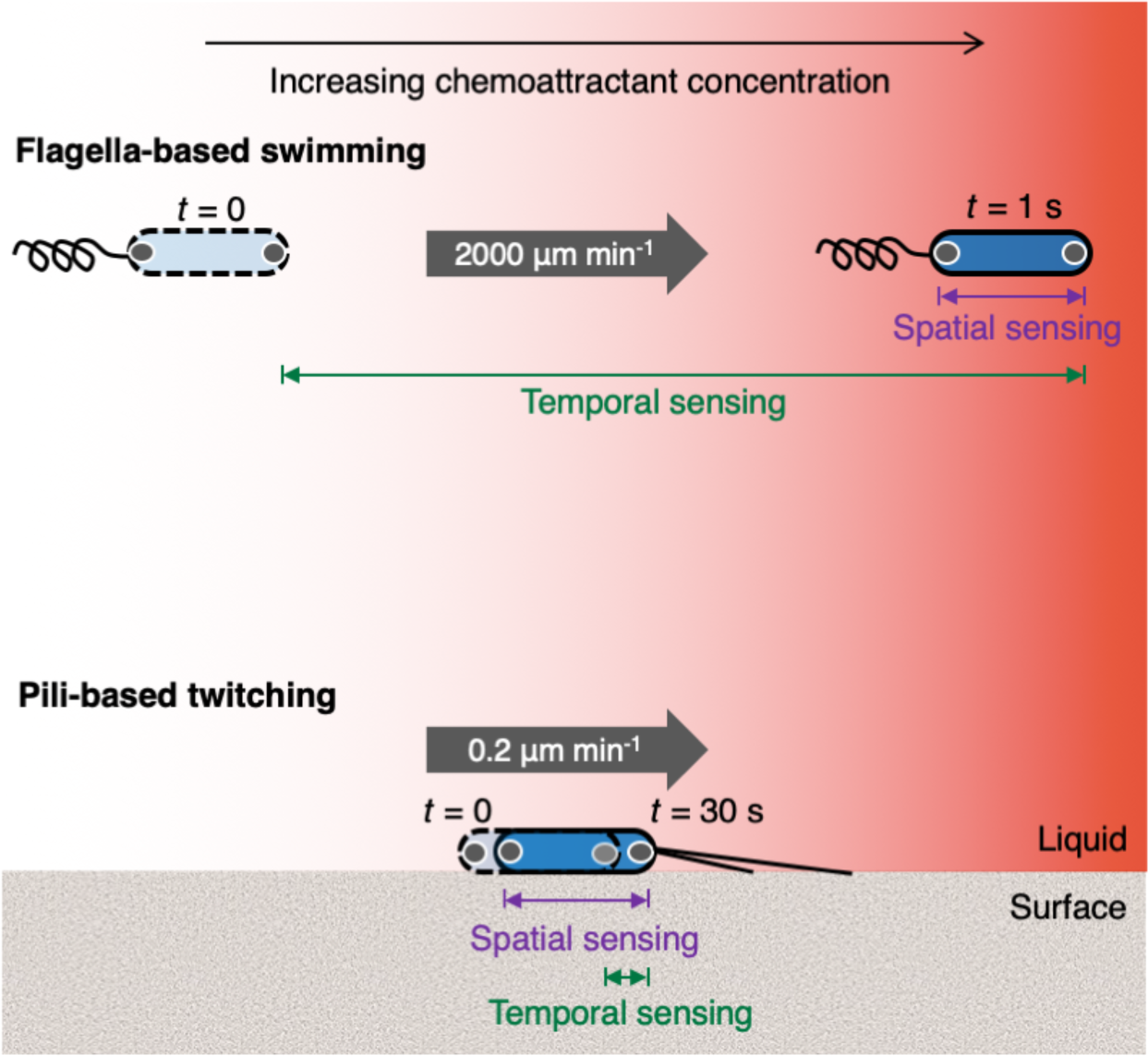
Swimming bacteria experience larger changes in concentration over time, whilst twitching bacteria experience larger changes in concentration over the lengths of their bodies. In principle, chemotaxing cells could either sense changes in chemoattractant concentration by moving from one location to another and comparing how the concentration changes over time (temporal sensing) or by directly comparing differences in concentration over the length of their bodies (spatial sensing). The rapid speed of swimming bacteria (see for example [38]) means that over the course of their typical response time (order 1 s), they would experience a larger change in concentration in time than space (denoted by the green and purple bars respectively). The opposite is true for surface-attached twitching bacteria, which move much more slowly (**Fig. S1*C***) and have response times on the order of 30 s [32]. Here, chemoreceptor clusters are represented by the grey circles within the cell poles.

Whilst eukaryotic cells are capable of both forms of sensing, the paradigm in the study of bacterial chemotaxis is one of temporal sensing. In particular, swimming bacteria have long been known to use temporal sensing [9–11] and their rapid motility allows them to measure changes in concentration that occur over length scales equivalent to tens of cell body lengths (**Fig. 1**). The molecular mechanisms that facilitate temporal sensing in swimming bacteria have been resolved in a number of different species and are particularly well understood in swimming *Escherichia coli* [12–16]. To the best of our knowledge, there is only one potential observation of spatial sensing in bacteria, which was suggested as an explanation for the U-shaped trajectories made by an uncultured bacterium collected from marine sediments that swims using flagella extending from each of its two poles [17]. However, these analyses were not definitive and the widely held belief in the literature is that spatial sensing is typically not feasible in swimming bacteria due to physical constraints posed by their small size and rapid movement [9-11, 18-21]. Consistent with this view, whenever the chemosensory systems of swimming bacteria have been characterised in detail, they have exclusively been found to use temporal sensing mechanisms to detect chemical gradients [22–28].

This focus on swimming cells contrasts with the fact that most bacteria live in surface-attached communities called biofilms [29–31]. Flagella are ineffective at driving motility in surface-attached cells [32–34]; instead they propel themselves using other forms of motility [35, 36]. For instance, many surface-attached bacteria move via twitching motility, which is driven by the extension and retraction of type IV pili that function like molecular grappling hooks to pull cells across surfaces [37]. It was previously demonstrated that individual *Pseudomonas aeruginosa* cells can use twitching motility to navigate chemoattractant gradients [32]. Specifically, when exposed to a chemoattractant gradient that alternated direction, surface-attached cells were observed to rapidly reverse direction in response, typically before traveling a single micron. In contrast to swimming cells that reverse direction by switching the direction of flagellar rotation [38], twitching cells reverse direction by switching pili activity to the opposite pole of their rod-shaped bodies [39, 40]. However, it is not known how surface-attached *P. aeruginosa* cells resolve which of their poles is directed toward higher chemoattractant concentrations as they navigate chemical gradients.

*A priori,* there are good reasons to suspect that surface-attached *P. aeruginosa* cells might use a different type of gradient sensing compared to swimming cells (**Fig. 1, Supplementary Information**). On average, twitching cells migrate approximately four orders of magnitude more slowly than swimming cells [32, 38]. However, on short timescales, twitching motility is true to its name and the dynamics of individual pili cause cells to jerk back and forth relative to chemical gradients as they travel [37, 41]. Therefore, if twitching cells were to use temporal sensing, they would have to resolve the relatively small and slow temporal changes in chemoattractant concentration that result from a cell’s overall movement direction, from the large, stochastic fluctuations that result from the cell’s jerking back and forth relative to the chemical gradient. One might thus expect that spatial sensing could particularly benefit twitching cells because it would allow them to decouple their unsteady motility from their ability to measure chemical gradients (**Fig. 1**). Furthermore, it is known that surface-attached bacteria are able to detect non-chemical stimuli, such as light and mechanical forces, over the lengths of their bodies [40, 42]. We therefore decided to investigate whether surface-attached *P. aeruginosa* cells, like eukaryotes, can detect chemical gradients across their cell bodies.

### Twitching bacteria do not use temporal gradients to guide chemotaxis

While one can argue how spatial sensing might benefit twitching *P. aeruginosa* cells (**Fig. 1, Supplementary Information**), the well-documented temporal mechanisms used by swimming cells suggest that temporal sensing is more likely. For this reason, we began by testing whether temporal changes in chemoattractant concentration could explain the directed motility of *P. aeruginosa* on surfaces. The experiments that documented pili-based chemotaxis used a dual-flow microfluidic device where molecular diffusion mixes two streams of fluid with different chemoattractant concentrations as they flow down the length of the device (**Fig. S1**, [32]). In these assays, cells undergoing chemotaxis simultaneously experience a spatial gradient over the length of their bodies as well as temporal changes in chemoattractant concentration as they move along the gradient. This makes it difficult to ascertain whether cells are responding to either spatial or temporal stimuli.

To directly test whether twitching cells use temporal signals to guide chemotaxis, we developed a custom microfluidic set-up that uses Taylor-Aris dispersion [43, 44] to generate a concentration gradient of succinate – a preferred carbon source of *P. aeruginosa* and a known chemoattractant [32] – that flows past cells. Importantly, this device exposes all cells to an approximately equal temporal stimulus, independent of their movement speed or direction (**Fig. 2, Methods**). Cells in dual-flow microfluidic experiments bias their motility by increasing or decreasing their reversal frequency when moving away from or towards chemoattractants, respectively (**Fig. S1**, [32]). Therefore, if cells indeed used temporal measurements to guide chemotaxis, we would expect that a temporal decrease in succinate concentration would cause the cells in our Taylor-Aris dispersion experiments to reverse more frequently, and vice versa.

**Figure 2.**
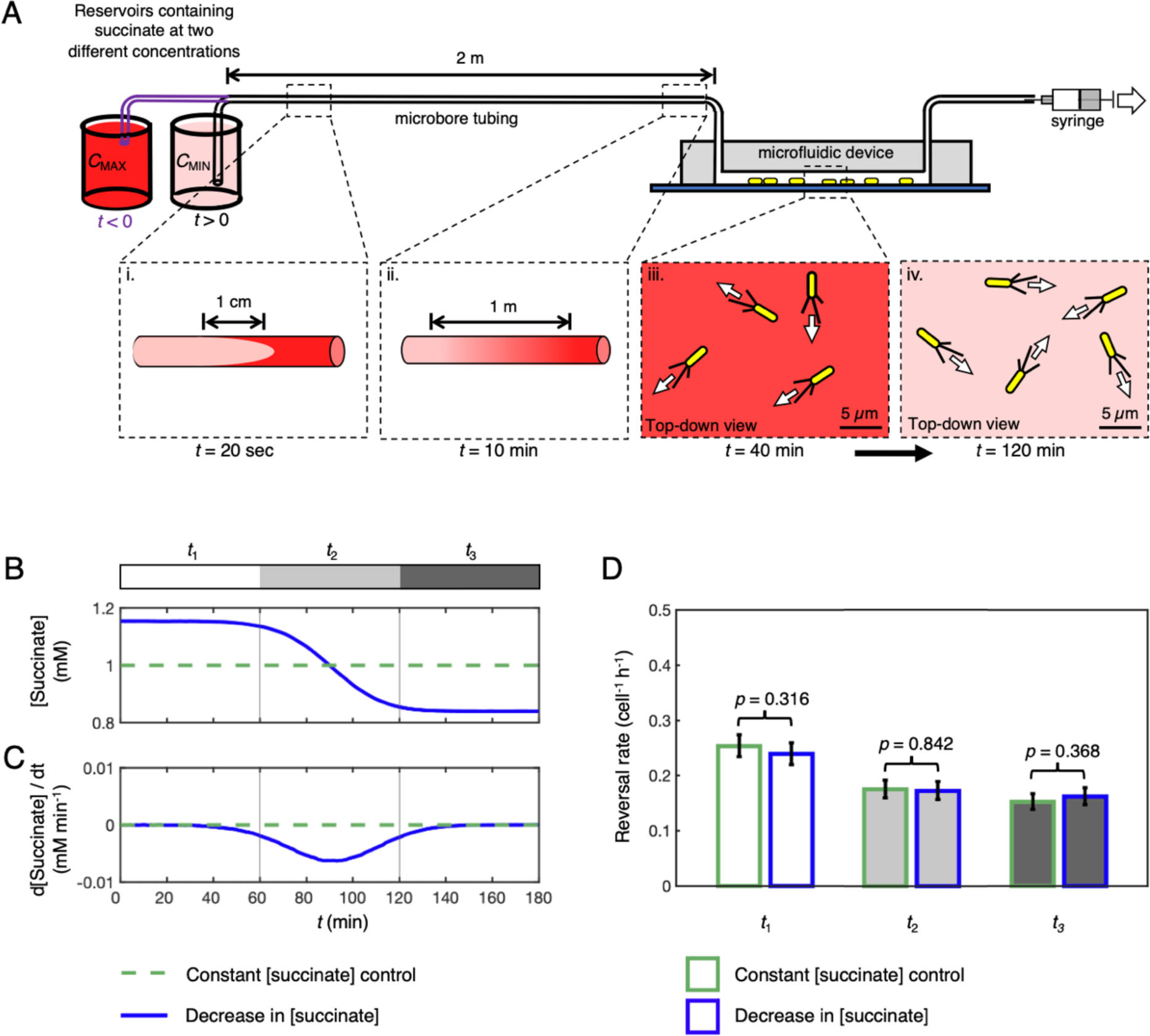
Temporal changes in concentration do not induce a chemotactic response in surface-attached *P. aeruginosa*. **(A)** To expose cells to a temporal concentration gradient with only a minimal spatial gradient, we used Taylor-Aris dispersion to generate a concentration gradient over the length of a 2 m long tube, which was then flowed past cells attached to the surface of a microfluidic device. To accomplish this, we first filled the entire system with media containing a concentration of *C*MAX = 1.16 mM of succinate. At *t =* 0, the upstream end of the tubing was transferred into a reservoir containing a lower concentration of succinate (*C*MIN = 0.84 mM) and fluid was pulled through the system using a syringe pump. Since fluid moves fastest through the tube along its centerline, a bullet shaped plug of lower concentration fluid forms within the tube (panel i) but is then rapidly mixed across the width of the tube via molecular diffusion (panel ii). By the time the interface between the two different fluids reaches the microfluidic device, it has formed a longitudinal gradient along the length of the device with a length-scale of approximately 1.6 m. Therefore, cells attached to the interior of the device experience a smooth decrease in concentration over time (panel iii – panel iv). **(B, C)** By labelling the fluid in the first reservoir with dye, we quantify the succinate concentration and temporal concentration gradient at each time point (blue lines; dashed green lines show a control with a constant succinate concentration *C* = 1 mM, **Methods**). This experiment exposes cells to approximately the same mean temporal concentration gradient that cells experienced in the dual-inlet experiments where chemotaxis was observed (**Fig. S1**, [32]), but with a ∼16,000-fold smaller spatial gradient. **(D)** Using automated reversal detection, we first confirmed that in the 1 h period before the succinate gradient entered the microfluidic device (time interval *t*1; white bar, blue outline), the reversal rate was statistically indistinguishable from the reversal rate during the same time period in a simultaneous control experiment where a constant concentration of succinate was maintained throughout (white bar, green outline). Specifically, a one-sided exact Poisson test (**Methods**) did not reject the null hypothesis that these two reversal rate measurements come from the same Poisson distribution, *p* = 0.316. Similarly, the reversal rates in the presence of a temporal succinate gradient (time interval *t*2; light grey bar, blue outline) and in the 1 h period after the gradient had cleared the microfluidic device (time interval *t*3; dark grey bar, blue outline) were statistically indistinguishable from the reversal rates during the same time periods in the control (*p* = 0.842 and *p* = 0.368). The total number of reversals observed in our six simultaneously imaged fields of view was *n*r = 1496 and 1391 across a total of *n*t = 468,596 and 439,632 trajectory points in the control and experimental conditions respectively. Error bars show 95% confidence intervals assuming that reversals follow a Poisson distribution (**Methods**). Data shown is representative of two bio-replicates (see **Fig. S4**).

We designed our Taylor-Aris dispersion experiments to expose cells to the same average chemical temporal stimuli that cells experienced in the dual-flow experiments where chemotaxis was originally demonstrated. This correspondence was accomplished by matching both the concentrations (*C*) and mean temporal concentration gradients (d*C*/d*t*) that cells experience in those experiments (see **Methods**). Importantly, in our Taylor-Aris dispersion experiments, the chemoattractant gradient forms over the length of a two-meter-long tube leading to the microfluidic device (**Fig. 2*A***), such that the chemical gradient measures approximately 1.6 m in length by the time it reaches the cells. In contrast, in dual-flow experiments, the gradient instead forms across the width of the microfluidic device and has a characteristic length-scale of 100 µm. Therefore, the cells in our Taylor-Aris dispersion experiments experience approximately a 16,000-fold smaller gradient across the length of their bodies (i.e. d*C*/d*x*) compared to the dual-inlet experiments, whilst experiencing approximately the same mean temporal stimuli (d*C*/d*t*).

We used massively parallel cell tracking and automated reversal detection [32] to quantify the movement of thousands of cells attached to the surface of a microfluidic device (**Fig. S2**). In addition to exposing cells to temporal gradients of succinate, we also ran a control experiment in an adjacent microfluidic channel on the same microscope where cells were exposed to a constant succinate concentration over time, allowing us to distinguish any potential changes in cell motility induced by the temporal succinate gradient from other, more general changes in cell motility that result from the physiological adaptation of cells to the surface [45] and increasing cell density on the surface caused by in-situ cell division [46] over the course of these ∼3 h long experiments (**Fig. S2**). First, to establish a baseline, we analysed cell motility in the one hour period that preceded the succinate gradient entering the microfluidic device (white region labelled *t*1, **Fig. 2*BC***) and compared it to that measured over the same time period in the control. As reversals are relatively rare events [32], we imaged six fields of view in each channel, which allowed us to track approximately 1 x 10^5^ cells simultaneously (see **Fig. S2**). We found that the baseline reversal rate prior to the gradient entering the microfluidic channel (white region labelled *t*1, **Fig. 2*B***) was statistically indistinguishable in both the experimental and control channels over this time period (**Fig. 2*D*, Fig. S3-5**). This strong correspondence thus indicates that we can directly compare the cellular reversal rates in the two channels at later time points to assess whether a temporal gradient in concentration causes cells to alter their reversal rate.

We next calculated the reversal rate of cells as they experienced a temporal decrease or increase in succinate concentration (light grey region labelled *t*2, **Fig. 2*B,C***) and compared it to that measured over the same time period in the constant succinate concentration control. Regardless of whether cells were exposed to a temporal increase or decrease in succinate concentration, cell reversal rates in time period *t*2 were statistically indistinguishable when compared between experimental and control conditions (**Fig. 2*D*, Fig. S3-5**). Finally, we measured reversal rates in the one hour time period after the temporal gradient had cleared the microfluidic device to confirm that the gradients did not have a latent effect on cell reversal rates (dark grey region labelled *t*3, **Fig. 2*B,C***). Once again, cell reversal rates in time period *t*3 were statistically indistinguishable when comparing between the control and experimental conditions (**Fig. 2*D*, Fig. S3-5**). Taken together, our results thus strongly suggest that cells do not alter their reversal rate in response to temporal succinate gradients. Whilst it is known that twitching cells generate chemotaxis by actively modulating their reversal frequency in response to the direction that they are travelling along a chemoattractant gradient (**Fig. S1**, [32]), the absence of a response in our Taylor-Aris dispersion experiments suggests that *P. aeruginosa* cells do not use the mean temporal changes in concentration they experience to guide pili-based chemotaxis.

However, we decided to explore another possible basis for temporal sensing. While the Taylor-Aris dispersion experiments simulated the long-term, average temporal changes in concentration experienced by cells in experiments where chemotaxis was observed, on shorter timescales, twitching cells routinely undergo much more rapid movement caused by the stochastic release of individual pili [37, 41]. These movements can momentarily transport a cell 1-2 orders of magnitude faster than its average speed and could potentially elicit a behavioural response by exposing a cell to larger temporal stimuli. This is because the magnitude of the temporal gradient a cell experiences scales with cell velocity, *V*, relative to a chemical gradient like d*C*/d*t* = *V* d*C*/d*x*. Therefore, to measure the response of twitching cells to more rapid changes in succinate concentration, we used a programmable microfluidic system that smoothly switches between two different concentrations of succinate over a period of 1.5 min, yielding temporal gradients, d*C*/d*t*, that are approximately 40-fold larger than the experiments shown in **Fig. 2*C***, (**Methods**). Given the short timescale of these temporal gradients, we alternated between two different succinate concentrations more than 12 times over the course of the experiment, allowing us to expose the same cells to both positive and negative temporal concentration gradients and analyse data across them separately. While these temporal gradients were much sharper than those in the Taylor-Aris dispersion experiments, we again found that temporal stimuli did not generate any detectable changes in cells’ reversal rate (**Fig. S6**). Taken together, these first experiments strongly suggest that chemotaxis in surface-attached *P. aeruginosa* is not guided by temporal stimuli alone.

### A sub-cellular reporter to quantify chemotactic behaviour in stationary cells

Our first experiments indicated that twitching chemotaxis is not driven by temporal sensing, suggesting instead that *P. aeruginosa* cells might directly sense differences in concentration across the length of their bodies. However, to evaluate this possibility, we needed to find a way to experimentally decouple the spatial and temporal information that cells experience. The challenge is that a cell moving through a steady spatial gradient of chemoattractant will experience differences in concentration along the length of its body, whilst simultaneously experiencing changes in concentration over time as it moves relative to the gradient. To decouple these two different stimuli from one another, we decided to study the behaviour of stationary cells, which typically make up a relatively small percentage of cells within our microfluidic assays (approximately 5-10%). The question then was how does one characterise chemotactic behaviour in cells that are not moving?

Here, we initially found inspiration in the studies of *Myxococcus xanthus*, which can also move via twitching motility [47]. Reversals occur 40 times more frequently in *M. xanthus* and are accompanied by changes in the sub-cellular localisation of two motor proteins, PilB and PilT, which are responsible for pili extension and retraction respectively [48–50]. In twitching *M. xanthus* cells, PilB localises to the front pole of a moving cell (the “leading pole”), whilst PilT localises predominantly to the rear pole (the “trailing pole”). The two motor proteins then switch between the two poles of *M. xanthus* cells during reversals. If these motor proteins exhibit similar patterns of localisation in twitching *P. aeruginosa* cells, we could potentially use fluorescent fusions to quantify reversals in cell polarity, even in cells that are temporarily stationary.

To visualise the retraction motor PilT in cells undergoing reversals, we fused PilT to yellow fluorescent protein (YFP) and expressed it in a *P. aeruginosa* strain lacking a functional native copy of PilT (*ΔpilT*::*pilT-yfp*, see **Methods**). This fusion protein fully complemented the motility of the *ΔpilT* strain (**Fig. S7**), a mutant lacking the first portion of the gene’s coding region (**Methods**, see [51]). We find that our PilT-YFP fusion protein localises predominantly to the leading cell pole in twitching *P. aeruginosa* cells (**Fig. 3*B*** and ***C***) and re-localises to a cell’s opposite pole during reversals (**Fig. 3*D***), which is consistent with two recent studies [40, 52]. Given that PilT instead localises to the trailing pole in twitching *M. xanthus* cells, this implies that different molecular mechanisms are used to generate reversals in these two species.

**Figure 3.**
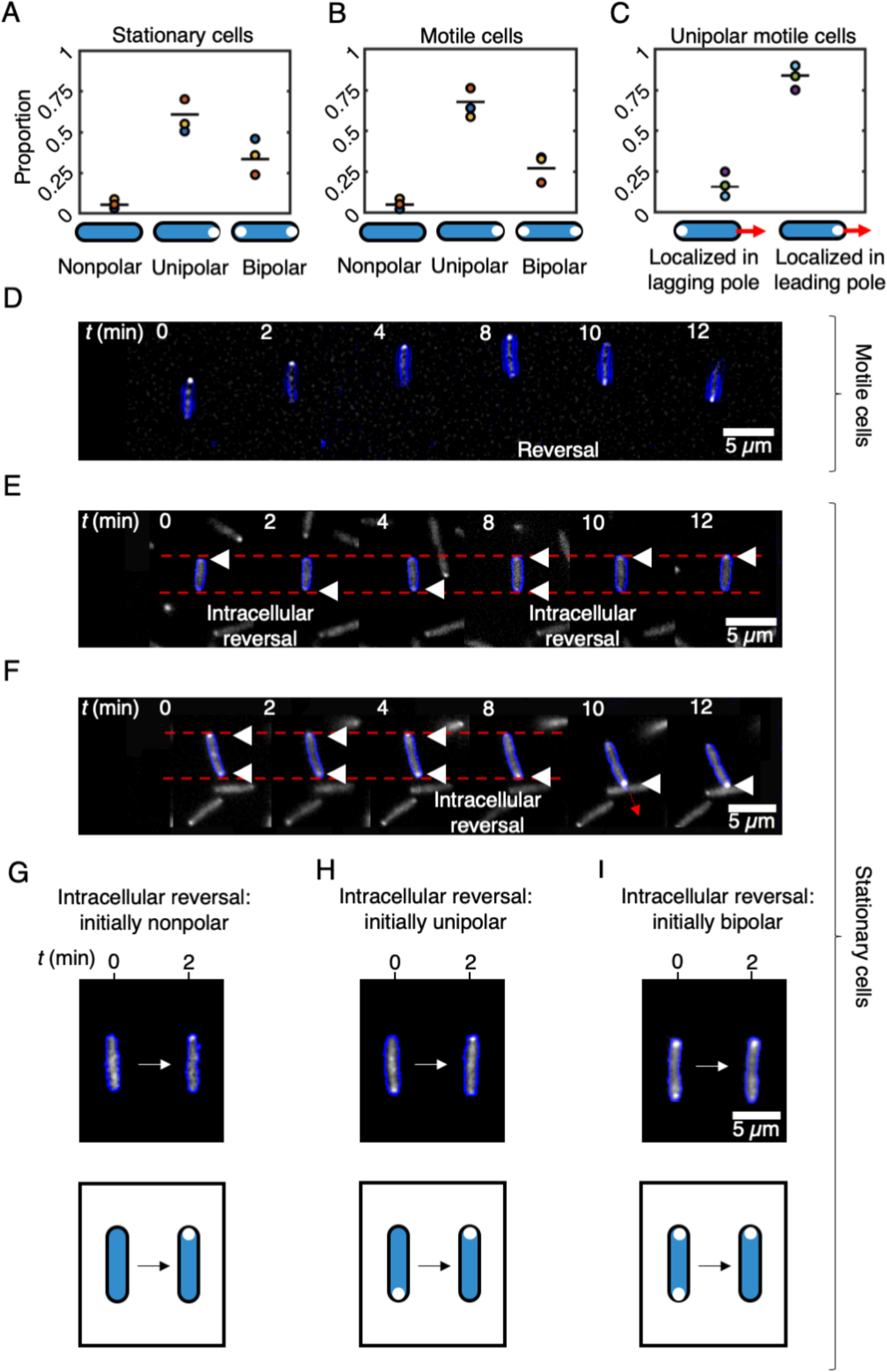
PilT-YFP localises to the leading pole of motile cells and can dynamically re-localise within the bodies of both motile and stationary cells, providing a means to infer chemotactic behaviour. **(A, B)** In the majority of both stationary (A), and motile cells (B), the PilT-YFP fusion protein localises to one of the two cell poles (unipolar). A smaller proportion of cells have PilT-YFP localisations in both poles (bipolar) or lack appreciable localisations altogether (nonpolar). Black lines show the mean of three bio-replicates that were each conducted on different days, represented here with a different coloured circle. The data from each bio-replicate contained over *n* = 1000 trajectories. **(C)** If we consider only those motile cells that have a unipolar PilT-YFP localisation, we find that PilT-YFP is significantly more likely to localise to a cell’s leading pole (a two-sided binomial test of proportions rejects the null hypothesis of equal proportions with *p* < 1 x 10^−18^). **(D)** A time series of a motile twitching cell (cell outline shown in blue) undergoing a reversal at *t* = 8 min. PilT-YFP (shown in white) localises to the leading pole, so that it swaps from one pole to the other when the cell reverses direction. **(E)** A time series of a stationary cell reveals that PilT-YFP can swap between a cell’s two poles over time, an event we call an “intracellular reversal”. Localisations of PilT-YFP are marked with white triangles. **(F)** A cell that is initially stationary has PilT-YFP localised to both of its poles, but subsequently PilT-YFP accumulates within its bottom pole shortly before the cell initiates movement in the downward direction. Faint dashed red lines in E and F mark the position of the two cell poles in the first image of the timeseries. **(G, H, I)** Intracellular reversals can occur in cells that are initially nonpolar (G), unipolar (H) or bipolar (I).

In stationary cells, PilT-YFP can also localise to either neither (“nonpolar”), one (“unipolar”) or to both cell poles simultaneously (“bipolar”, Fig. 3*A*). Crucially, we found that the localisation of PilT-YFP remains dynamic in the stationary cells in our microfluidic assays, with new localisations forming and old localisations dissipating over time (**Fig. 3*E*** and ***F***). These findings indicate that changes in the sub-cellular localisation of PilT-YFP can be used to distinguish between the leading and lagging pole before a cell starts to move. Specifically, this fusion allows us to detect “intracellular reversals” in stationary cells, which occur when PilT-YFP redistributes within the cell (**Fig. 3*G-I***) and quantify how they are elicited by different types of chemical gradients. Tracking changes in the sub-cellular localisation of PilT-YFP therefore allows us to analyse the chemotactic behaviour of stationary cells.

### *P. aeruginosa* uses spatial sensing to guide chemotaxis across surfaces

To test for spatial sensing, we used a custom Y-shaped microfluidic device [32] to expose our *P. aeruginosa* (*ΔpilT*::*pilT-yfp*) cells to a spatial gradient of succinate that alternates in direction (**Fig. 4*A***). We then followed the distribution of PilT-YFP within >1000 stationary cells to quantify how they respond to a gradient that alternated direction approximately every 45 min (**Methods**). Stationary cells that underwent intracellular reversals can be separated into two different categories: “correct” intracellular reversals in which cells re-localise PilT-YFP towards the pole experiencing higher succinate concentrations and “incorrect” intracellular reversals, where PilT-YFP is re-localised towards the pole experiencing lower succinate concentrations (Fig. 4*B*).

**Figure 4.**
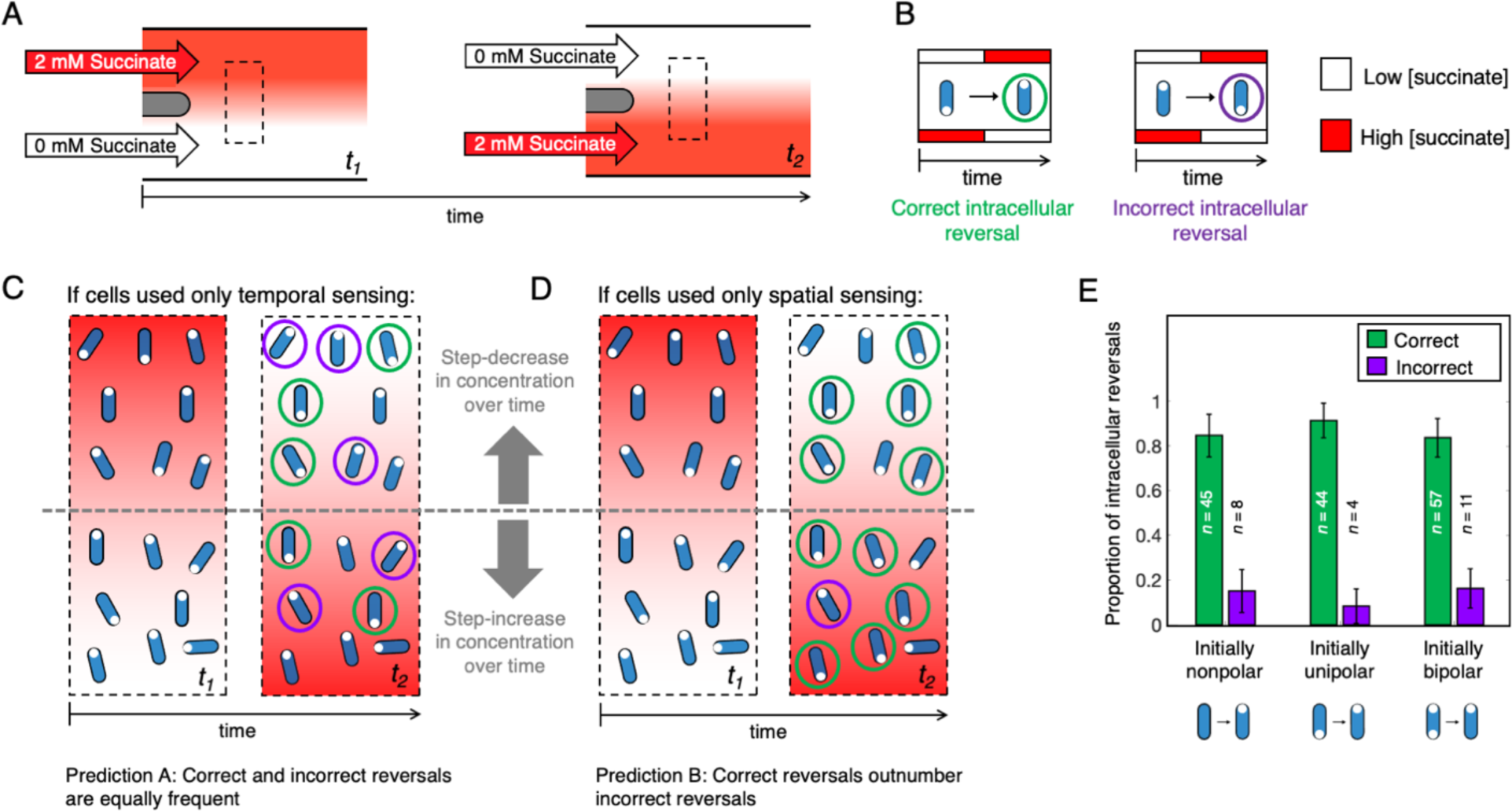
Intracellular reversals in stationary cells exposed to an alternating succinate gradient preferentially re-localise PilT-YFP to the cell pole experiencing larger succinate concentrations, indicating that they are capable of spatial sensing. **(A)** We used a dual-flow microfluidic device to expose cells to a spatial gradient of succinate that alternates direction [32]. The dashed black box indicates the region downstream of the two inlets, where we imaged cells. **(B)** In response to this alternating spatial gradient, stationary cells (blue) expressing PilT-YFP (white circles) can undergo either correct or incorrect intracellular reversals. **(C, D)** The relative proportion of correct and incorrect intracellular reversals in this experiment can be used to determine whether cells use spatial or temporal sensing. (C) Stationary cells using only temporal sensing could garner no information about a gradient’s spatial orientation and would therefore be equally likely to generate correct and incorrect intracellular reversals (“Prediction A”). (D) In contrast, stationary cells capable of spatial sensing could directly sense the gradient’s spatial orientation, allowing them to deploy correct intracellular reversals at a greater frequency than incorrect intracellular reversals (“Prediction B”). **(E)** Quantifying the behaviour of 171 stationary cells undergoing intracellular reversals within our alternating gradient experiments (see **Appendix** and **Supplementary File 1**) revealed that correct intracellular reversals occurred approximately 6 times more frequently than incorrect intracellular reversals, regardless of whether PilT-YFP localization was initially nonpolar, unipolar or bipolar (see **Fig. 3*G-I***). An exact two-tailed binomial test rejected the null hypothesis that correct and incorrect intracellular reversals were equally abundant with *p* = 2.37 x 10^−7^, 1.51 x 10^−9^ and 1.28 x 10^−8^ for nonpolar, unipolar and bipolar intracellular reversals respectively. This is consistent with Prediction B, indicating that cells are capable of directly sensing differences in concentration over the length of their bodies. Error bars show 95% confidence intervals.

The relative frequency of correct and incorrect intracellular reversals in stationary cells allows us to directly test whether cells respond to temporal or spatial stimuli. Since stationary cells do not move appreciably relative to the gradient, the temporal stimuli they experience do not encode information that could allow them to determine the orientation of the chemical gradient. Instead, on one side of the device stationary cells simply experience an increase in concentration over time, whilst on the other side, they experience a decrease in concentration over time (**Fig. 4*C*** and ***D***). Therefore, temporal sensing and spatial sensing lead to two different, and easily distinguishable, predictions in these experiments. If stationary cells used temporal sensing, intracellular reversals would be independent from the gradient’s orientation, so one would expect that correct and incorrect intracellular reversals would both occur randomly and, therefore, at approximately the same rate (“Prediction A”, **Fig. 4*C***). In contrast, if stationary cells can make spatial measurements, we expect that correct reversals will occur more often than incorrect reversals. This is because cells that sense the direction of the chemical gradient by directly measuring it across their bodies would be able to correctly ascertain the gradient’s spatial orientation (“Prediction B”, **Fig. 4*D***).

Across three bio-replicates, we identified 171 cells that were stationary following the change in gradient orientation and subsequently undertook intracellular reversals (**Fig. 4*E***; see **Appendix** and **Supplementary File 1**; a detailed description of how intracellular reversals were identified is given in the **Methods**). A fraction of stationary cells sometimes began to move off after the gradient changed direction before observably altering their PilT-YFP distribution, so we also used cell movement to diagnose the chemotactic response of these initially stationary cells (**Methods**). Separating these 171 intracellular reversals by direction revealed a striking result: correct intracellular reversals occurred approximately 6 times more frequently than incorrect ones (148 correct, 23 incorrect; **Fig. 4*E***), suggesting therefore that twitching cells directly sense chemoattractant gradients across the length of their cell bodies. This trend is remarkably consistent across stationary cells regardless of whether their initial PilT-YFP localisation is nonpolar, unipolar, or bipolar (**Fig. 4*E***). Moreover, cells were observed to correctly determine the direction of the succinate gradient despite being subjected to sharp changes in succinate concentration over time (**Fig. 5*A-C*, S8**; **Appendix** and **Supplementary File 1**). These temporal changes in concentration were 2-3 orders of magnitude larger than those in the Taylor-Aris dispersion experiments, indicating that spatial sensing is robust to large temporal changes in concentration, such as the random fluctuations that arise from twitching cell’s jerky movement relative to a chemical gradient. Lastly, we note that twitching *P. aeruginosa* cells always exhibit a basal level of reversals even in the absence of chemical gradients [32], which means that a proportion of incorrect intracellular reversals are expected, albeit at a lower frequency than correct ones (**Fig. 4*D,E*, Fig. 5*D***).

**Figure 5.**
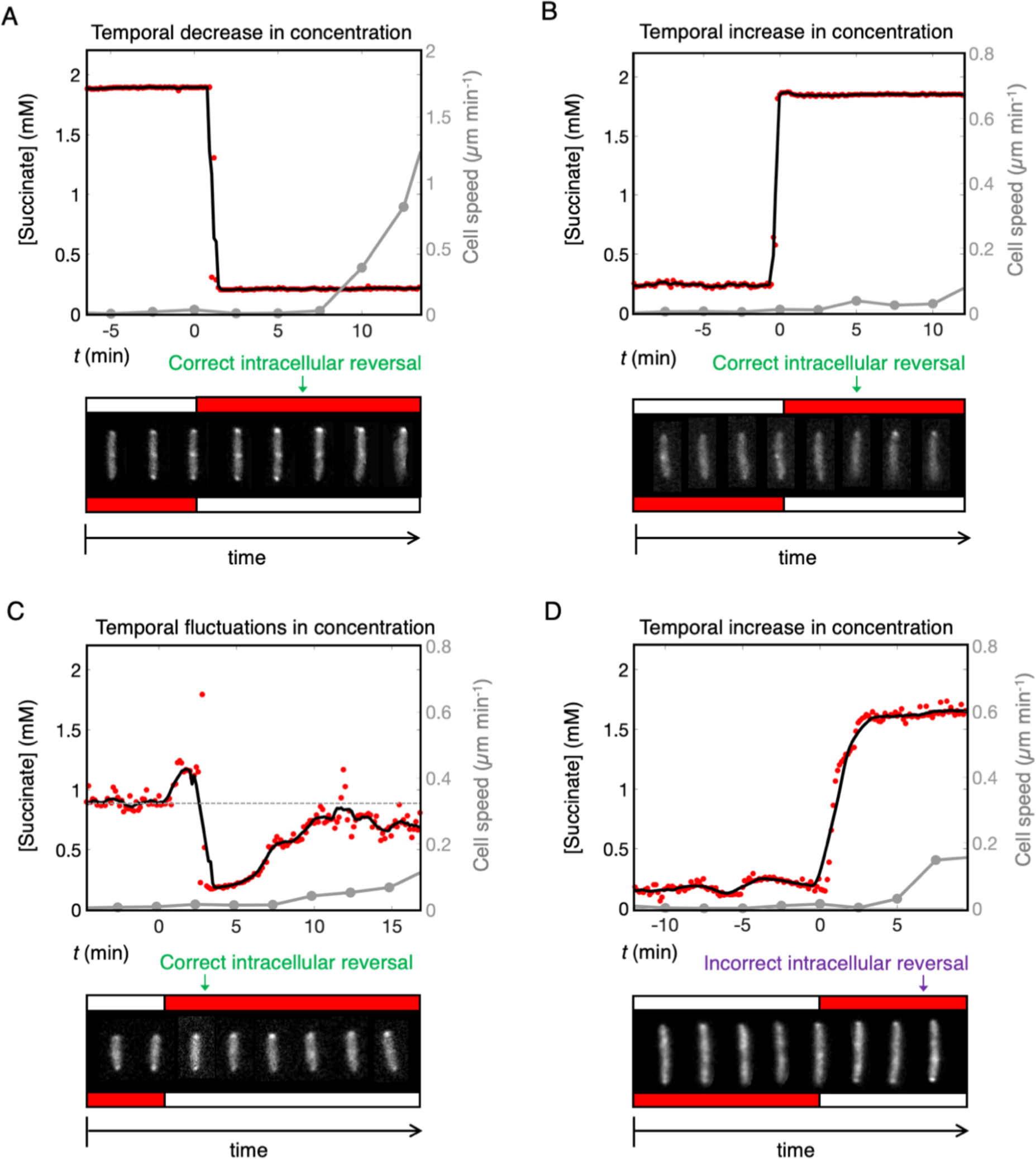
Stationary cells can sense changes in the orientation of a chemoattractant gradient, despite large temporal fluctuations in concentration. We simultaneously quantified the succinate concentration that a cell experienced over time (red circles, black line shows moving average), cell speed (grey line), and the intracellular distribution of PilT-YFP, as cells were exposed to a succinate gradient that alternates direction. Grey circles indicate the timepoints for which cell images are shown (i.e. at 2.5 min intervals). To help guide the eye, cells shown in the time-series at the bottom of each plot have all been repositioned so that they are vertically oriented and their centroid remains at a fixed position. **(A)** This cell experiences a sharp temporal decrease in succinate concentration when the gradient changes direction. PilT-YFP is observed to re-localise to the cell pole that is now exposed to the higher chemoattractant concentration (a correct intracellular reversal), and the cell later moves off in the direction of its new leading pole. PilT-YFP is shown in the bottom inset, with red and white boxes indicating the direction of high and low succinate concentration respectively. **(B)** A second example shows a cell experiencing a sharp increase in succinate concentration over time and also undergoing a correct intracellular reversal. While PilT-YFP fusion protein is initially nonpolar in this cell, it subsequently re-localises exclusively to the cell pole positioned in a higher succinate concentration. **(C)** A third example shows a cell that was positioned close to the centreline of the succinate gradient such that when the gradient alternated direction, it experienced noisy fluctuations in succinate concentration, including both increases and decreases in concentration. Despite this, the cell also underwent a correct intracellular reversal – PilT-YFP fusion protein was initially localised to both poles (with no observable directional polarity) and it subsequently re-localised exclusively to the cell pole positioned in a higher succinate concentration. **(D)** Although less frequent, cells were also observed to undergo incorrect intracellular reversals. Here a cell experiencing an increase in succinate concentration over time relocalised PilT-YFP to the cell pole positioned in a lower succinate concentration and subsequently moves in that direction. While these four examples of intracellular reversals are representative, movies of each of the intracellular reversals that we observed can be found in the **Appendix**, along with a description of how each was classified in **Supplementary File 1**.

The temporal changes did produce interesting trends, however. We observed more intracellular reversals in cells experiencing a decrease in succinate concentration over time compared to cells experiencing an increase (**Fig. S8**). This result is consistent with previous studies that find a cell’s response to a chemical gradient depends on both the gradient’s strength and the absolute concentration it experiences [53, 54]. Yet still, correct reversals outnumbered incorrect reversals in both cases, with correct reversals outnumbering incorrect ones by approximately 10-fold when the concentration was decreasing, whilst an approximately 4-fold difference was observed when the concentration was increasing (**Fig. S8**). These results suggest that cells can correctly identify the direction of the spatial gradient across the lengths of their bodies regardless of the sign of the temporal gradient. Taken together, our data therefore show that *P. aeruginosa* cells robustly navigate chemoattractant gradients using spatial sensing.

## Conclusion

We find that surface-attached *P. aeruginosa* cells can directly measure differences in concentration over the length of their bodies. In contrast, the signal transduction systems that guide chemotaxis in diverse swimming bacteria, including *P. aeruginosa*, use temporal sensing [9–11]. The use of spatial sensing was previously thought to be confined to the sophisticated signal transduction systems of eukaryotic cells [8, 18]. Eukaryotic spatial sensing is regulated by a molecular “compass” composed of intracellular chemical gradients. These gradients are generated from competition between rapid excitatory signalling generated by chemoeffector-chemoreceptor binding and slower, cell-wide inhibitory signalling, known as localised excitation, global inhibition or LEGI interactions [55, 56]. Bacterial cells are typically an order of magnitude smaller than eukaryotic cells and as a result, diffusion is predicted to smooth out any intracellular protein gradients within the bacterial cytoplasm approximately one hundred times more rapidly [57, 58]. Despite this limitation, bacteria have been shown to establish cytoplasmic gradients of protein phosphorylation by localising the proteins driving phosphorylation and de-phosphorylation to opposite cell poles [19]. Furthermore, it has recently been demonstrated that twitching *P. aeruginosa* cells are able to sense differences in mechanical stimuli across the lengths of their bodies via the two response regulators of the Pil-Chp chemotaxis-like system (PilG and PilH, which may prove comparable to the eukaryotic-like LEGI system [40]). As well as sensing mechanical stimuli, the Pil-Chp system is also thought to play a role in twitching chemotaxis [32] and it is therefore possible that similar LEGI interactions could facilitate spatial measurements of chemical gradients in *P. aeruginosa*.

Bacteria commonly live on surfaces, where they form biofilms. Our results show that the well-established paradigm of bacterial chemotaxis, based on measuring changes in concentration over time, does not hold for surface-based movement in *P. aeruginosa*. Instead, we find that cells navigate on surfaces using spatial information. This mode of sensing is well suited to the slow movement and steep chemical gradients associated with biofilm life and, relative to temporal sensing, it likely would allow twitching cells to measure larger changes in concentration, enhancing their ability to discriminate chemical gradients from stochastic noise (Fig. 1, **Supplementary Information,** [20, 59]). Indeed, our experiments demonstrate that even stationary cells can use spatial information to sense chemical gradients. This observation raises the possibility that static bacteria living in mature biofilms could use gradient sensing to guide biofilm development.

## Methods

### Bacterial strains and culturing

Wild-type *P. aeruginosa* PAO1 (Kolter collection, ZK2019) was used as the model organism for this study. To visualise the localisation of PilT within cells, we sought to express a fluorescently labelled copy of PilT from the native promoter of *pilT* on the chromosome. However, we were not able to detect any fusion protein using this approach with epi-fluorescent imaging, presumably because the native expression levels of *pilT* were too low. We therefore sought an alternative solution. Firstly, we generated a *pilT* mutant lacking the first portion of the gene’s coding region in our model PAO1 strain using a previously published plasmid kindly gifted to us for this study (pJB203, [51]; we refer to this mutant as *ΔpilT*). We then generated a PilT-YFP protein fusion expressed from a low-expression promoter (BG35) previously characterised in *Pseudomonas putida* [60]. Briefly, *pilT* was amplified from the chromosome of PAO1 using two primers that were complementary to the sequence immediately downstream of the *pilT* start codon (PILT_F) and ≈100 base pairs downstream of the *pilT* stop codon (PILT_R, see **Table 1** for primer sequences). The coding sequence of YFP was amplified from the plasmid pEYFP-N1 (Clontech) using an upstream primer (YFP_F) that additionally introduced the BG35 promoter immediately upstream of a ribosome binding site (designed using automated methodology described by [61]) and a downstream primer (YFP_R) that introduced a rigid linker [62] to separate the functional domains of the two amplified proteins (YFP and PilT). These two amplified fragments were then combined by secondary PCR, ligated into the linearized vector pGEM-T (Promega) and transformed via electroporation into *E. coli* S17-1, a broad-host range donor strain. We then used a previously established protocol for using a mini-Tn7 system to insert our *pilT-yfp* construct into the chromosome of our *ΔpilT* strain at its single *att*Tn7 site ([63], *Δ*(*pilT*)*att*Tn7::*pilT-yfp*). Doing so restored the motility of our *ΔpilT* strain to WT levels, thus confirming that our PilT-YFP fusion protein is functional when expressed from the BG35 promoter at the chromosomal *att*Tn7 site (**Fig. S7**). The final construct was confirmed by sequencing.

**Table 1.**
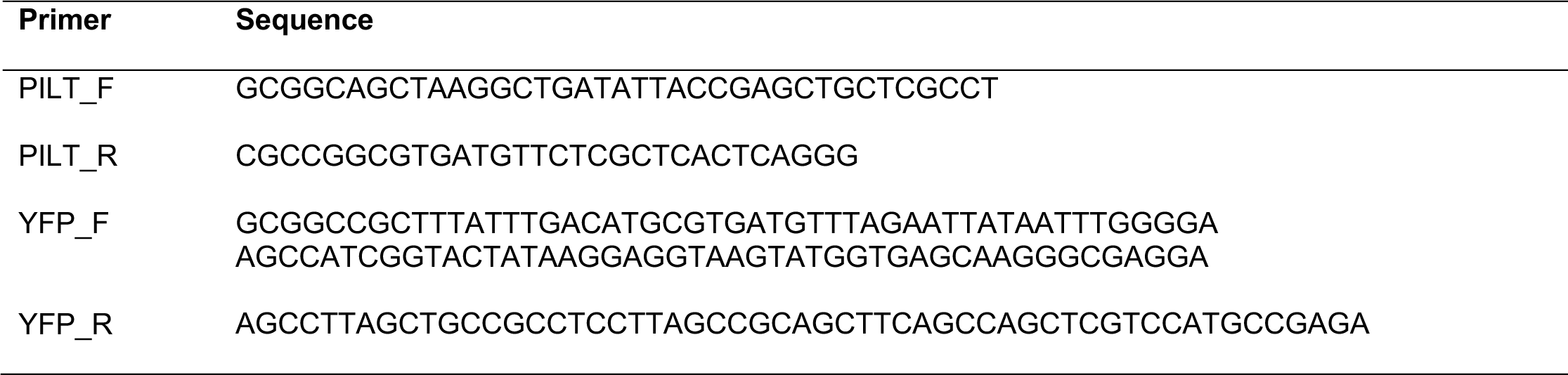
Sequences of primers used in this study.

All strains were grown from frozen stocks overnight in LB (Fisher, 37°C, 250 rpm) and sub-cultured (1:30 dilution) in tryptone broth (TB, 10 g L^−1^, Bacto tryptone) for 2.5 h to obtain cells in exponential phase. Cells were then diluted to an optical density at 600 nm of either 0.15 (experiment shown in **Fig. 2** and **S2-S5**) or 0.5 (all other experiments) in TB media before being used to inoculate microfluidic experiments.

### Imaging

In the Taylor-Aris dispersion experiments (**Fig. 2, S2, S3, S4, S5** and **S6**), we used a Nikon Ti2-E inverted microscope equipped with a “Perfect Focus” system and a Hamamatsu Orca-Fusion camera. In the experiments that quantified the distribution of PilT-YFP (**Fig. 3*A-C*, 4, 5** and **S8**) and for the experiment shown in **Fig. S1**, we used a Nikon Ti-E inverted microscope equipped with a “Perfect Focus” system, a Hamamatsu Flash 4.0 v2 camera and a CoolLED pE-4000 illuminator. For the experiment shown in **Fig. S7**, we used a Zeiss Axio Observer inverted microscope equipped with a “Definite Focus” system, a Zeiss AxioCam MRm camera, and a Zeiss HXP 120 illuminator. We used 20X Plan Apochromat air objectives throughout, except for our studies of the subcellular localisation of our PilT-YFP fusion protein, which used a 60X Plan Apochromat oil-immersion objective (on the Nikon Ti-E system).

### Microfluidic experiments

Our custom-designed devices were cast with PDMS (Sylgard 184, Dow Corning) using molds fabricated from SU-8 on silicon wafers (FlowJEM, Toronto, Canada). Holes for tubing were punched through the PDMS using a Harris Unicore 1.5 mm biopsy tool (Agar Scientific). The PDMS was then bonded to glass coverslips (50 mm by 75 mm, No. 1.5 thickness, Agar Scientific) using a corona treater (BD-20AC, Electro-Technic Products), as previously described [64].

We plumbed the inlets and outlets of our microfluidic devices using Tygon microbore tubing (1.5 mm outside diameter) and then placed the entire setup in a vacuum chamber for 1 h to reduce the potential for air bubbles. The devices were then mounted onto the microscope and the outlet tubing was connected to a 10 ml plastic syringe (Luer-Lok, Becton Dickinson) using a 23-gauge needle (PrecisionGlide, Becton Dickinson). The syringe was filled with nutrient media (TB) and mounted onto a syringe pump (PhD Ultra, Harvard Apparatus). To remove air from the system, we first injected TB through the device at a flow rate of 100 µl min^−1^. Exponential-phase cells (as described above) were then drawn into the device via suction at a flow rate of 50 µl min^−1^ through the inlet tubing. Once cells reached the test section of the channel, all inlets and outlets were clamped using hemostats for 10 min, which allowed cells to attach in the absence of any flow. After this attachment period, the TB from the syringe was injected through the device at 100 µl min^−1^ for 10 min to remove any remaining planktonic cells. Lastly, the ends of the inlet tubing were placed into new reservoirs and fluid was pulled through the device via suction for the remainder of the experiments.

The experiments shown in **Fig. S1** and **S7** were performed using the commercial BioFlux 200 microfluidic system (Fluxion Biosciences), using protocols that have been previously described [32].

### Taylor-Aris dispersion microfluidic experiments to expose cells to temporal chemical gradients

For the experiments shown in **Fig. 2** and **S2-S5**, we used a custom microfluidic device with a single inlet and outlet at either end of a rectangular microfluidic channel (30 mm in length with a cross section 1 mm wide and 75 µm deep). The inlet was connected to a 2 m length of Tygon tubing whose other end was placed in a reservoir containing TB mixed with succinate and the entire system was filled with this fluid. Subsequently, we moved the end of the tube to another reservoir, containing a different concentration of succinate. When this new fluid was drawn into the tube via suction, Taylor-Aris dispersion mixed the interface between the media containing the two different concentrations of succinate longitudinally along the length of the 2 m tube before it flowed over the top of the cells. Alternatively, for control experiments, the end of the inlet tube was inserted into reservoirs that both contained TB with 1 mM succinate. Thus, cells in these control experiments did not experience any chemical gradients.

As discussed in the main text, our Taylor-Aris dispersion experiments were designed to expose cells to approximately the same mean concentration (*C*) and temporal concentration gradient (d*C*/d*t*) that cells experienced in the dual-flow experiments where pili-based chemotaxis towards succinate was readily observed (**Fig. S1**, [32]). In these experiments, the static spatial gradient of succinate had a magnitude of approximately d*C*/d*x* = 0.02 mM μm^−1^. Individual twitching cells moved along this gradient with average speed of *V*C *=* 0.2 μm min^−1^ (see **Fig. S1*C***) and thus experienced a temporal gradient of succinate on the order of d*C*/d*t* = *V*C d*C*/d*x* = (0.2 μm min^−1^). (0.02 mM μm^−1^) = 0.004 mM min^−1^. Cells in this region of the device experienced an absolute concentration of succinate of *C* ≈ 1 mM.

Compared to flagella-based swimming, the motility of surface-attached *P. aeruginosa* cells is relatively slow and reversals are relatively rare – on average, a cell reverses direction only once every several hours [32]. To ensure that our results were statistically robust, we aimed to collect as many cell trajectories (and thus reversals) as possible over the course of an experiment. To achieve this, we first used an automated microscope stage to simultaneously image six different fields of view within each microfluidic channel every minute (a total of twelve different scenes as we imaged in two channels simultaneously). Second, we aimed to expose cells to a temporal change in succinate concentration that lasted a period of approximately one hour, so that we could detect a sufficient number of reversals over this period (labelled *t*2 in **Fig. 2*B-D*, S3** and **4**).

The length-scale of the succinate gradient that forms along the length of the inlet tube is set by competition between molecular diffusion in the radial direction and differential advection in the longitudinal direction of the tube, such that the length-scale of the gradient in the tube increases with the flow rate. To obtain succinate gradients with the correct magnitude, we used previously described theory [43] to design our experimental procedure. We first inserted the end of the inlet tube into the reservoir containing succinate at the higher concentration, *C*MAX, and then filled the entire microfluidic system with this media via suction. Then we switched the inlet tube to the reservoir containing the lower succinate concentration, *C*MIN, and pulled this second media into the inlet tube at a rate of 20 μl min^−1^ for 10 min. This formed a succinate gradient within the tube leading to the microfluidic device. We then lowered the flow rate on our syringe pump to 2 μl min^−1^ allowing us to extend the timescale of the temporal gradient that subsequently passed over the cells within the device.

We observed that the succinate gradient took approximately τ = 60 min to pass through the microfluidic channel, as visualised by using dye (Chicago Sky Blue 6B, 0.5 mg ml^−1^) in each run of the experiment (E.g., **Fig. 2*B***). This dye does not affect pili-based chemotaxis in *P. aeruginosa* [32] and is predicted to have approximately the same distribution as the succinate given that they both have a similar molecular weight. We chose *C*MAX = 1.16 mM and *C*MIN = 0.84 mM, which yielded a d*C*/d*t* ≈ (*C*MAX – *C*MIN) / τ = (1.16 mM – 0.84 mM) / 60 min = 0.005 mM min^−1^ and ensured that cells experienced an average concentration of 1 mM succinate over the course of the experiment, which also matched the uniform succinate concentration used in control experiments. Our Taylor-Aris dispersion experiments thus closely matched the mean temporal gradient and mean concentration of succinate observed in the previously described dual-flow experiments (d*C*/d*t* ≈ 0.004 mM min^−1^and *C* ≈ 1 mM, respectively).

The cells in our Taylor-Aris dispersion experiment primarily experience temporal variations in concentration that result from the spatial gradient of succinate flowing past them. We note that the speed of cells in our experiment *V*C = 0.2 μm min^−1^ (see **Fig. S1*C***) is orders of magnitude smaller than the speed at which the succinate gradient passes through the device (approximately 27,000 μm min^−1^), so a cell’s movement relative to the gradient has no appreciable impact on the temporal variation in succinate concentration they experience. Moreover, the length-scale of the succinate gradient when it passes through the test section of the microfluidic device is approximately *L =* (27,000 μm min^−1^). (60 min) = 1.6 m. Thus, the spatial gradient of succinate that cells experience across the length of their bodies in the Taylor-Aris dispersion experiments can be estimated as d*C*/d*x* ≈ (*C*MAX – *C*MIN) / *L* = (1.16 mM – 0.84 mM) / 1.6 m = 2.0 × 10^−7^ mM µm^−1^, which is several orders of magnitude smaller than the spatial gradients that cells experienced in the dual-flow experiments (d*C*/d*x* ≈ 0.02 mM μm^−1^).

In summary, the cells in the Taylor-Aris dispersion experiments experience approximately the same mean temporal stimuli as they do in the previous dual-flow experiments, whilst experiencing spatial gradients that are only vanishingly small in comparison.

To follow cell motility in these experiments, images were captured in brightfield at a rate of 1 frame min^−1^. Using Fiji [65], we stabilised the timeseries of brightfield images using the Image Stabiliser plugin to remove drift in the *x*, *y* plane. Next, the background was made more homogenous using the Normalise Local Contrast plugin and the intensity of the background was reduced using the Subtract Background feature. Finally, a bleach correction plugin was used to correct for long-term changes in the relative pixel intensity of the cells in brightfield compared to the background, which varies as the concentration of dye changes over time [66]. Cells were then tracked using the Trackmate plugin for Fiji [67]. Finally, to analyse cell motility and to detect when cells reverse direction, we used an image analysis pipeline in Matlab that we developed previously to study twitching motility in *P. aeruginosa* [32].

### Experiments to test whether cells can respond to sharp temporal variations in concentration owing to pili release events

Twitching motility is characteristically jerky and cells frequently undergo rapid displacements caused by the release of single pili, causing them to briefly move 1-2 orders of magnitude faster than their average speed [37, 41]. We thus tested the possibility that twitching cells in the presence of chemical gradients might employ a temporal sensing modality that is tuned to respond to these relatively short but steep temporal chemoattractant gradients.

For these experiments, we used a dual-inlet BioFlux 200 microfluidic system (Fluxion Biosciences) in which one inlet was connected to TB mixed with a larger concentration of succinate (*C*MAX = 1.16 mM), while the other inlet was connected to TB mixed with a smaller concentration of succinate (*C*MIN = 0.84 mM). Instead of passing fluid through both inlets simultaneously so they formed a spatial gradient within the test section [32], we instead passed fluid through only one inlet at a time, which exposes all cells in the test section to the same succinate concentration. We used computer-controlled software to alternate the flow between the two inlets, such that cells sequentially experienced a rapid increase in succinate concentration followed by a rapid decrease in succinate concentration over time. Like the Taylor-Aris dispersion experiments described in the previous section, we chose these *C*MAX and *C*MIN values so that the mean succinate concentration that cells experienced was 1 mM, which was the concentration where chemotaxis was observed to peak in the dual-flow experiment where cells where exposed to a spatial gradient of succinate.

We added Chicago Sky Blue 6B dye (0.5 mg ml^−1^) to the media containing the higher concentration of succinate (*C*MAX), whilst the media containing the lower concentration of succinate (*C*MIN) did not contain dye. By quantifying the change in dye intensity at the downstream end of the test section of the device, we observed that cells experienced a smooth change in concentration between the two different media over a timescale of τ ≈ 1.5 min (**Fig. S6**). Because the time period of the temporal gradient (τ) in these experiments is relatively short and therefore affords less time to observe reversals, we alternated the flow between the two inlets every 15 min so that we could expose cells to at least six increases and decreases in succinate concentration over the course of one experiment (**Fig. S6*a***). We observe that the transition between the two succinate concentrations occurs smoothly and consistently in the test section of the device. We note that the overall duration of our microfluidic experiments is limited because in-situ cell division eventually crowds the surface, which makes tracking individual cells difficult.

We can estimate the temporal gradient in these experiments as d*C*/d*t* ≈ (*C*MAX – *C*MIN)/*τ* = (1.16 mM - 0.84 mM) / 1.5 min = 0.2 mM min^−1^, **Fig. S6*B*** and ***C***), which is 1-2 orders of magnitude larger than the temporal gradients that cells were exposed to in the Taylor-Aris dispersion experiments described in the previous section and is approximately the same strength as the temporal stimuli that we predict a cell in our dual-flow experiments will experience momentarily during pili release events [37, 41, 68]. We can estimate the spatial gradients that form over the length of the test section in these experiments as d*C*/d*x* ≈ (*C*MAX – *C*MIN) / (*U* τ) = (1.16 mM – 0.84 mM) / (2500 µm min^−1^. 1.5 min) = 8.5 × 10^−5^ mM µm^−1^, (where *U* is the mean flow speed), which is approximately 200-fold smaller than the spatial gradients that cells experienced in the dual-flow experiments (d*C*/d*x* ≈ 0.02 mM μm^−1^).

To follow cell motility, two fields-of-view were imaged in brightfield at a higher frame rate of 7.5 frames min^−1^. Using Fiji [65], images were processed and tracked using the Trackmate plugin [67] as described above. To analyse cell motility and to detect when cells reverse direction, we once again used our previously developed image analysis pipeline in Matlab [32].

To test whether cells can sense and respond to this larger temporal stimulus, we compared cells’ reversal rate before, during, and after they experienced a temporal gradient in succinate concentration across six increases and six decreases in succinate concentration (**Fig. S6*B*** and ***D***). Our statistical analyses found that neither an increase nor a decrease in succinate concentration elicited cells to change their reversal rate (**Fig. S6*C*** and ***E***). These experiments thus show that surface-attached *P. aeruginosa* cells do not respond to the larger temporal gradients that they would experience during pili release events.

### Quantifying the sub-cellular localisation of our PilT-YFP fusion protein

To measure how the localisation of our PilT-YFP fusion protein varies from a cell’s leading pole to its lagging pole, we developed an image analysis pipeline that automatically tracks cell position, length, and orientation in brightfield and uses this information to quantify the distribution of YFP using the corresponding epi-fluorescence images. Brightfield images were captured at a frame rate of 7.5 frames min^−1^, while epi-fluorescence images to visualise YFP were simultaneously acquired at a lower frame rate of 0.5 frames min^−1^. The higher frame rate for brightfield allowed us to track cell motility with sufficient accuracy, whilst the lower frame rate for the YFP imaging allowed us to avoid bleaching and phototoxicity.

All preliminary image analysis was conducted in Fiji [65]. Brightfield images were processed as outlined above. Epi-fluorescence images were processed in the same way as brightfield images, except we additionally used a Difference of Gaussian filter to enhance the contrast of the localised accumulations of PilT-YFP.

The cells in these processed images were then tracked using software called the Feature-Assisted Segmenter/Tracker (FAST, [68]; see also https://mackdurham.group.shef.ac.uk/FAST_DokuWiki/ <u>dokuwiki/doku.php?id=start</u>) which allowed us to track cell position and orientation with greater precision compared to the tracking plugins available in Fiji. To map how the distribution of PilT-YFP varies along the cell length and how that distribution changes as cells move, we used the cell centroid, length and orientation extracted from brightfield images tracked using FAST to resolve the region corresponding to each cell on the respective YFP epi-fluorescence image. To accurately quantify the distribution of the fusion protein, we needed to develop a method that could detect PilT-YFP localisations even when they were slightly offset from the cell’s centreline, in addition to being robust to small amounts of cell movement that occurred in the time interval between when the brightfield and YFP images were captured. To account for these factors, for each cell we calculated the YFP fluorescence intensity along a series of parallel lines with the same orientation and length as the cell, but separated by a small distance from one another so that collectively they spanned a width slightly larger than the cell’s width. We then recorded the maximum YFP intensity that occurred across all of these lines, to obtain the maximum fluorescence intensity at each position along the cell’s length. This process was used to record the distribution of PilT-YFP in the several thousand images of individual cells that were recorded across three bio-replicates. We omitted from our analyses any YFP intensity values that were below a minimum threshold (corresponding to the background YFP intensity observed outside of cells) to prevent the small number of mis-tracked cells from influencing our results. We also omitted any cells with an aspect ratio smaller than 1.4, which ensured that our analyses only included cells that were attached to the surface by both cell poles.

We next quantified the distribution of PilT-YFP fusion protein within the poles of the cells. Because the maximum YFP intensity often does not occur at the very tip of the pole, we measured the maximum YFP intensity in the vicinity of the poles. The cell length was measured using YFP images and the maximum YFP intensity was calculated in the two regions at either end of the cell, each corresponding to 1/10 of the overall cell length. To classify the distribution of PilT-YFP within a cell as nonpolar, unipolar, or bipolar (**Fig. 3**), we normalised the maximum YFP intensity within each pole by the mean YFP intensity in the central ¼ of the cell. If the normalised YFP intensity in a given pole (denoted as *I*1 and *I*2 for pole 1 and pole 2) exceeded a threshold *I*MIN (determined by visual inspection for each bio-replicate) the protein was considered to have aggregated within that pole. More specifically, if both *I*1 > *I*MIN and *I*2 > *I*MIN the cell was considered bipolar, whereas if either *I*1 > *I*MIN or *I*2 > *I*MIN the cell was considered unipolar. Lastly, if *I*1 < *I*MIN and *I*2 < *I*MIN the cell was considered nonpolar.

To increase the accuracy of the automated assignment of cells as bipolar, unipolar, or nonpolar, we also implemented the following two rules:

- When *P. aeruginosa* nears cell division, the pili machinery (and thus the PilT-YFP protein fusion) begins to localise additionally to the nascent cell poles, which are positioned at mid-cell [69]. In such instances, the maximum fluorescence intensity can occur in the mid-cell region rather than at the poles. As we are interested in the processes underlying cell motility (rather than cell division), we excluded cells from our analyses whose average fluorescence in the middle ¼ of their bodies was larger than that found in either of the poles.
- Cells initially assigned as being bipolar were re-assigned as unipolar if *I*1 and *I*2 differed by a fixed ratio (determined by visual inspection for each bio-replicate). This allowed us to ensure that we assigned cells with strongly asymmetrical patterns of PilT-YFP localisation as “unipolar”, rather than “bipolar”.

To compare the distribution of PilT-YFP in stationary and moving cells, we classified trajectories by their speed (**Fig. 3**). Due to pixel noise and the effect of fluid flow, the measured trajectories of non-motile cells exhibited a finite velocity. To account for these effects, we classified cells moving slower than 0.038 μm min^−1^ as “stationary”, whilst cells moving faster than this threshold were classified as “motile”. To prevent cells simply jostling back and forth from being considered motile, we additionally removed trajectories from the motile category whose net to gross displacement ratio (NGDR) was less than 0.04. In addition, we excluded cells that were actively rotating from the motile category, by identifying cells whose bodies had an angular velocity larger than 0.073 radians min^−1^ for a contiguous period of longer than 2 min. These angular velocities were obtained from measurements of cell orientation that had been smoothed with a first order Savitzky-Golay filter (using a 20 min window) to reduce noise. All the parameters used in these analyses were extensively ground-truthed to ensure that they had the desired effect.

### Generating alternating spatial chemoattractant gradients in custom microfluidic devices

To expose cells to a spatial chemoattractant gradient that alternates in direction by 180 degrees, we used a custom microfluidic device described in detail previously [32]. Briefly, the device is composed of a Y-shaped channel with four inlets (two inlets in each branch of the Y) and a single outlet that was connected to a syringe pump.

In these experiments, a steady spatial gradient of succinate forms along the centreline of the device, where the fluids from two inlets located in opposite arms of the Y-shaped channel meet one another. The fluid supplied through one arm contained nutrient media (TB) supplemented with 2 mM of succinate and Chicago Sky Blue 6B dye (0.5 mg ml^−1^), whilst fluid from the other arm contained only undyed nutrient media. Molecular diffusion generated a stable gradient of succinate across the width of the channel, which could be readily quantified by imaging the dye since they have a similar diffusion coefficient.

The syringe pump pulled media through the device via suction (5 μl min^−1^) from reservoirs connected to the four inlets of the device. A hemostat was used to clamp the tubing connecting two of the inlets at any given time. To change the direction of the gradient, the hemostat is removed from one pair of tubes and transferred to the other pair, which contain the same two fluids but in the opposite orientation (see [32] for details).

Brightfield images were captured at a frame rate of 7.5 frames per minute so that changes in the gradient and cell movement could be tracked at a high temporal resolution. Epifluorescence images of the cells were captured at a slower frame rate of 0.4 frames per minute to avoid bleaching of the PilT-YFP fusion protein and to prevent phototoxicity. The details of how cells were tracked and how the distribution of the fusion protein inside them was quantified is outlined below.

### Analysing the localisation of our PilT-YFP fusion protein in stationary cells exposed to an alternating spatial chemoattractant gradient

To directly test whether surface-attached *P. aeruginosa* cells are capable of spatial sensing, we exposed our *ΔpilTatt*Tn7::*pilT-yfp* strain to a spatial gradient of succinate that alternated direction using the microfluidic device outlined in the previous section. To exclude the possibility that cells could use temporal sensing to determine the orientation of the new succinate gradient, we only considered intracellular reversals that occurred in stationary cells (see main text). Because the PilT-YFP protein fusion tends to localise to a cell’s leading pole (**Fig. 3*C***), each intracellular reversal can be categorised according to whether the new leading pole of a stationary cell is oriented towards (“correct”) or away from (“incorrect”) increasing succinate concentrations following the change in gradient orientation (**Fig. 4**). In addition, we also classified intracellular reversals according to whether PilT-YFP was initially localised in both poles (bipolar), in only one pole (unipolar), or in neither pole (nonpolar) before the intracellular reversal occurred (see **Fig. 3*G-I***).

While our other analyses used automated cell tracking to quantify cell behaviour, we decided to detect and classify these intracellular reversals manually for two main reasons. Firstly, a relatively small number of intracellular reversals are observed in these experiments, so we wanted to follow the behaviour of every single cell and rigorously ground-truth all putative intracellular reversals to confirm that they were not erroneous. Secondly, many stationary cells reside in densely-packed groups, which help to stifle movement. However, densely packed cells are challenging to track using automated methods without occasional errors and it is difficult to measure an individual cell’s PilT-YFP distribution without inadvertently having it contaminated by the YFP signal produced by neighbouring cells. (Note that in other experiments that were analysed using automated cell tracking, we developed filters to specifically exclude cells that were clustered together.)

We analysed the behaviour of every cell that was visible in the 16 different fields of view collected over the course of three separate microfluidic experiments (**Appendix**) and classified them with a detailed set of rules (see below). Out of >1000 cells that were investigated, we identified 171 stationary cells that subsequently performed an intracellular reversal as defined by our rules. To prevent potential errors, a preliminary list of intracellular reversals was independently assessed by two co-authors (JHRW and WMD) and any discrepancies were reconciled before our final analyses. All 171 intracellular reversals are labelled in the supplementary movies that accompany this manuscript (**Appendix**), along with the details of how each was classified (**Supplementary File 1**).

Below we describe in detail the rules that were used to define and classify each putative intracellular reversal:

#### Identifying when an intracellular reversal occurs

We search for potential intracellular reversals in cells that are stationary after the succinate gradient changes direction. In many cases, stationary cells first localise PilT-YFP exclusively to their new leading pole before moving off, however, sometimes cell movement occurs first. An intracellular reversal therefore occurs as soon as a stationary cell either: (A) develops a unipolar pattern of PilT-YFP localisation that is different to that of its initial localisation of PilT-YFP or (B) moves off in a direction different to that of its initial localisation of PilT-YFP. In the first case, (A), a cell must re-localise PilT-YFP to a single pole in at least two of four consecutive frames (10 min) whilst in the second case, (B), a cell must move off in a consistent direction for at least two frames at a speed corresponding to at least one cell width per frame.

Following an intracellular reversal, we define a cell’s “new leading pole” as the one that either contains the new unipolar PilT-YFP localisation or leads its initial movement, whichever has occurred first. The orientation of a cell’s “new leading pole” after the intracellular reversal is then used to determine whether it can be classified as a “correct” or “incorrect” intracellular reversal by comparing its orientation relative to that of the new succinate gradient (**Fig. 4*A, B*** and ***C*** and see **Supplementary File 1**).

Importantly, for an intracellular reversal to have occurred, a cell must not have previously had a unipolar PilT-YFP localisation in the “new leading pole” in either: two or more of the four frames (10 min) that precede the appearance of the new succinate gradient or within the frame that immediately precedes the appearance of the new succinate gradient. This requirement thus ensures that cells have actively changed their distribution of PilT-YFP following the change in gradient direction and also prevents short-lived, random fluctuations in the distribution of PilT-YFP from being erroneously classified as an intracellular reversal.

#### Defining which cells are considered “stationary”

These experiments aim to analyse the behaviour of stationary cells because motile cells could potentially use temporal sensing to determine the orientation of the new succinate gradient. However, cells can sometimes exhibit small amounts of movement that are unrelated to their motility. For example, the flow in our experiments tends to push cells downstream whilst cells at the periphery of densely packed cell clusters can get pushed radially outwards by their neighbours as the cluster grows. As such movements are not under the active control of a cell, they could not encode information about the direction of a gradient via temporal changes in succinate concentration in the same way that active motility would. In addition, in our experiments, cells that are pushed a small distance by flow tend to move in the direction orthogonal to the gradient and thus do not experience appreciable changes in succinate concentration over time. We therefore monitor a cell’s movement in the direction along the gradient to ensure it is sufficiently small in the period preceding an intracellular reversal.

To determine whether a cell can be considered stationary, we monitor its movement from the frame after the last frame in which the initial succinate gradient was present until the frame in which the cell undergoes an intracellular reversal. However, since some intracellular reversals occur shortly after the gradient has changed direction, we also monitor cell movement for at least three frames (7.5 min) prior to any putative intracellular reversal. A cell is then considered “stationary” within these time periods provided that its centroid neither: (A) moves more than half a cell width in the same direction between two consecutive frames nor (B) moves more than one cell width at any point. All distances are measured along the direction of the chemical gradient and a cell width is approximately 0.9 µm.

Note that many cells are stationary for a finite period and so a cell that is currently stationary will likely have moved at some point in the past. Our analyses include cells that move whilst the initial succinate gradient is still present, but subsequently stop moving before the gradient starts to change direction. This is because such prior movement could not inform a cell that the orientation of the succinate gradient will change later in the experiment.

#### Assigning a cell’s polarity prior to an intracellular reversal

We categorise intracellular reversals according to the PilT-YFP localisation that they previously exhibited (**Fig. 3*G-I*** and **4*E***). For an intracellular reversal to be assigned as either nonpolar, unipolar or bipolar, the cell must have had that polarity mode more frequently than any other in the four frames (10 min) preceding the appearance of the final gradient orientation. If two different polarity modes are each present for two frames apiece, then we assign the polarity mode that occurs in the frame immediately preceding the appearance of the final gradient orientation. The “initial polarity” of 2 cells could not be resolved in these experiments because one of their cell poles was initially in very close proximity to that of their neighbours. The initial polarity of these cells was classified as “not assignable” in the **Appendix** and **Supplementary File 1**.

We also observed a small number (*n* = 13) of intracellular reversals in newly divided cells. If a cell that is stationary (as defined above) divides shortly after the change in gradient orientation, one or both of the resulting daughter cells could in theory undergo an intracellular reversal (as defined above). In these cases, the distribution of PilT-YFP is assigned as nonpolar, unipolar or bipolar (**Fig. 3*G-I*** and **4*E***) according to the most frequent localisation pattern in the frames between the cell division event and the subsequent intracellular reversal. We did not consider PilT-YFP localisations at the midpoint of the mother cell prior to cell division in our analyses, because they are not necessarily related to motility and can be asymmetrically divided between the two daughter cells during septation [69].

#### Assigning the temporal change in succinate concentration accompanying each intracellular reversal

As above, we used Chicago Sky Blue 6B dye to visualise the alternating succinate gradient. When the gradient changes orientation by 180 degrees, cells initially situated in regions of low succinate concentration (*C* < *C*MAX / 2, as determined by the dye intensity) experience a temporal increase in succinate concentration, whilst those initially in regions of high succinate concentration (*C* > *C*MAX / 2) experience a temporal decrease in succinate concentration. By following changes in the dye intensity, we were able to group intracellular reversals according to whether they occurred in cells that had experienced an overall increase or decrease in succinate concentration (**Fig. S8**). However, it was very difficult to distinguish the small temporal changes in succinate concentration (and thus dye intensity) experienced by cells situated close to the centreline of the spatial gradient. These cells (*n* = 10) were therefore excluded from the analyses that compared the response of cells experiencing a step-up in succinate concentration to a step-down in succinate concentration. The “temporal change in [succinate]” of these cells is marked as “not assignable” in the **Appendix** and **Supplementary File 1**.

### Methods used to illustrate the different types of intracellular reversals

We used automated cell tracking software (Trackmate plugin, Fiji, [65, 67]) to follow cell movement across four exemplar intracellular reversals in order to quantify changes in cell speed and to map changes in succinate concentration at the location of each of the four cells (**Fig. 5*A-D***). To ensure that we could obtain trajectories that spanned the entire length of experiment, the cells of interest were cropped out frame-by-frame using the “Brush Tool” in Fiji. This left us with only a single cell visible in the entire time-series of images, ensuring the automated tracking was not influenced by the presence of neighbouring cells. We used the resulting curated trajectories to calculate the projection of the cell’s velocity along the chemoattractant gradient (grey lines, **Fig. 5*A-D***). The concentration of succinate that a cell experienced over time (black lines in **Fig. 5*A*-*D***) was quantified using the Chicago Sky Blue 6B dye, which was mixed with the 2 mM succinate solution. The distribution of dye was imaged using brightfield microscopy and separate experiments demonstrated a linear dependence between the pixel intensity and dye concentration, allowing us to easily estimate the succinate concentration at the position of each cell within the device.

### Statistical analyses

To test whether cells use temporal chemoattractant gradients to guide pili-based motility, we developed statistical methods to determine whether cells alter their reversal rate in response to temporal gradients of succinate in comparison to control conditions where the concentrations of succinate were constant. Our Taylor-Aris dispersion experiments (**Fig. 2, S3-S5**) are ≈ 3 h long and the total number of cells changes over this time-scale due to cell detachment from and attachment to the surface, as well as continued cell division (**Fig. S2**, [46]). Furthermore, even in the absence of any chemical gradients, reversal rates change over time, likely driven by cells undergoing physiological adaptation following surface attachment ([45], see **Fig. S5**). To take these temporal trends into account, we divided our datasets into three time-bins corresponding to before, during and after the cells experienced a temporal gradient of succinate (see *t*1, *t*2, and *t*3 in **Fig. 2, S3** and **4**).

Our Taylor-Aris dispersion experiments imaged six different fields-of-view simultaneously at a frame rate of 1 frame min^−1^, yielding several thousand trajectories at each timepoint (**Fig. S2*A*** and ***C***). However, reversals are relatively rare - on average a cell reverses direction only once every several hours. Our datasets therefore consist of a very large number of time points at each of which a cell can either carry on moving in a relatively straight line or, with a low probability, reverse direction. We therefore assume that reversals are Poisson distributed, allowing us to calculate the confidence intervals of our reversal rate estimates. Using this assumption, we also used the exact Poisson test (using the “poisson.test” function in R) to test for differences in reversal rates between control and experimental conditions (**Fig. 2, S3** and **4**).

A similar approach was used to generate confidence intervals for our estimates of reversal rates for cells moving either up or down spatial chemoattractant gradients (**Fig. S7**). However, in these analyses, we calculated the mean reversal rate using data from the entire experiment (rather than subdividing it into different bins in time), because in these experiments, the gradient was present for the entire duration.

## Acknowledgements

We thank Joanne Engel and Yuki Inclan for strains and plasmids, Diego Gonzalez for help with cloning, Leslie Vanderpant for help designing microscope incubation chambers, Judy Armitage for advice, Oliver Meacock for help with automated cell tracking, and Sean Booth, Oliver Meacock and Matthias Koch for providing feedback on a previous version of this manuscript. This work was funded by a BBSRC DTP studentship (BB/J014427/1) awarded to JHRW; the Human Frontier Science Program (LT001181/2011L and RGY0080/2021), EPSRC Pump Priming Award (EP/M027430/1) and BBSRC New Investigator Grant (BB/R018383/1) to WMD; and by European Research Council Grant 787932 and Welcome Trust Investigator award 209397/Z/17/Z to KRF.

## Author Contributions

J.H.R.W., K.R.F., and W.M.D. designed research; J.H.R.W. performed research; J.H.R.W. and W.M.D. analyzed data; J.H.R.W., K.R.F., and W.M.D. wrote the paper.

## Data availability

The full data set that supports the findings of this study are available from the corresponding authors upon request. Supplementary movies are available to view at the following location: https://drive.google.com/drive/folders/10-SksdsZGDRTYGf6qQQi43qlcLZLk-Ee?usp=sharing

## Code availability

All of the code used to generate the findings of this study is available from the corresponding authors upon request.

## Strain availability

The bacterial strains used in this study are available from the corresponding authors upon request.

## Supplementary Information

### Planktonic and surface-attached bacteria face fundamentally different constraints when sensing chemical gradients

A number of arguments have been proposed to explain why swimming bacteria benefit from temporal sensing. Chief among them is that swimming bacteria typically move very rapidly, allowing them to travel tens of body lengths along a gradient before being reoriented by Brownian rotational diffusion [20, 70]. Thus, temporal sensing allows swimming bacteria to measure much larger changes in concentration over time than they could measure over the lengths of their bodies (**Fig. 1**). The bodies of twitching cells are not appreciably affected by Brownian motion as they are surface-attached and move four orders of magnitude more slowly (*V*C ≈ 0.2 μm min^−1^, see **Fig. S1*C***) compared to swimming cells (*V*C ≈ 2000 μm min^−1^ [38]). A twitching cell therefore travels the length of its body (mean ≈ 5 µm) in approximately 25 min. Thus, unless twitching bacteria have a very long memory, allowing them to measure changes in their chemical environment over periods longer than 25 min, spatial sensing across the cell body would allow cells to measure larger changes in concentration. In comparison, the temporal sensing systems used by swimming bacteria typically measure changes in concentration that occur over only a few seconds [70].

Furthermore, twitching cells tend to jerk back and forth as they move, owing to the stochastic detachment of individual pili [37, 41]. If twitching cells used temporal gradients to guide chemotaxis, they would therefore have to integrate a rapidly fluctuating signal (that frequently changes signs) over long timescales to ascertain whether they were moving up or down a chemical gradient. In contrast, spatial sensing would allow the cell to directly measure concentration differences between its two poles and so does not rely on cell movement. Indeed, we find that even stationary cells can sense chemical gradients.

These considerations indicate that planktonic and surface-attached bacteria may have evolved fundamentally different sensing mechanisms to sense chemical gradients.

## Supplementary Figures

**Figure S1.**
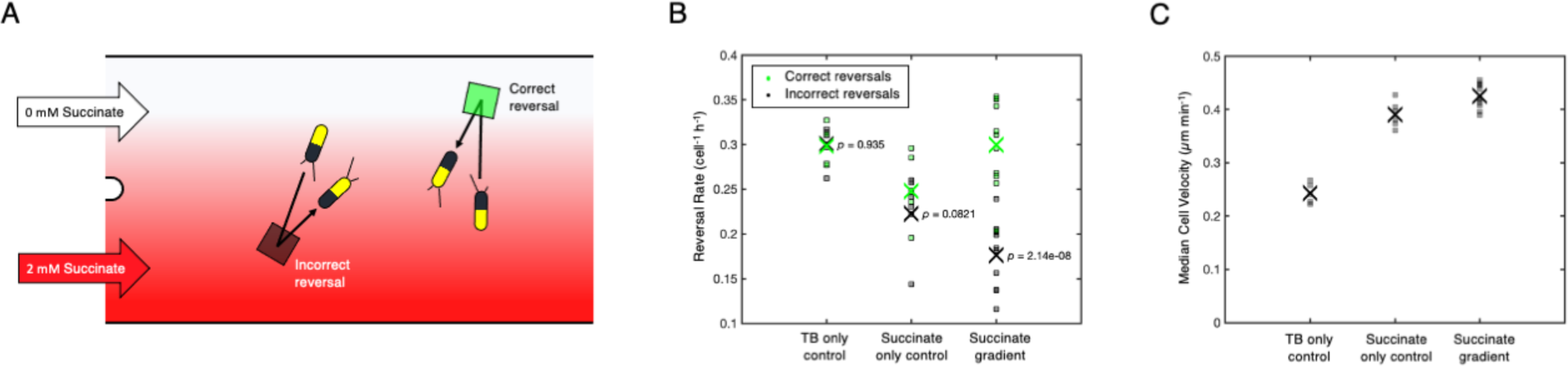
Surface-attached *P. aeruginosa* cells climb spatial succinate gradients by actively changing the rate at which they reverse direction. **(A)** A dual-inlet BioFlux microfluidic device was used to expose cells to a steady spatial gradient of succinate (*C*MIN = 0 mM, *C*MAX = 2 mM) with a characteristic length-scale of 100 µm, and an automated algorithm was used to detect when cells reversed the direction of their movement [32]. Reversals were classified as either “correct” or “incorrect”. Correct reversals (green square) occur in cells that were initially moving away from the source of succinate, whilst incorrect reversals (black square) occur in cells that were initially moving towards the source of succinate. **(B)** In the presence of a succinate gradient, the rate of correct reversals (green squares) is significantly greater than that of incorrect reversals (black squares), which drives chemotaxis of the population towards the source of succinate. The crosses (“X”) mark the mean of six separate TB-only control experiments, six separate succinate-only control experiments, or twelve separate succinate gradient experiments. In TB-only experiments, TB is passed through both inlets at the same time, whilst in succinate-only experiments, media containing succinate is passed through both inlets at the same time. These control experiments were processed in the same way as the experiment that used the succinate gradient, but since no gradient was actually present, the “correct” and “incorrect” rates shown for these controls are arbitrary. The *p*-values shown were obtained from paired *t*-tests, using the null hypothesis that the measured incorrect and correct reversal rates come from the same distribution. **(C)** Cell velocity is significantly higher in the presence of succinate gradients compared to both control experiments and is significantly higher in succinate-only controls compared to TB-only controls (one-way ANOVA, *p* = 2.22 x 10^−11^).

**Figure S2.**
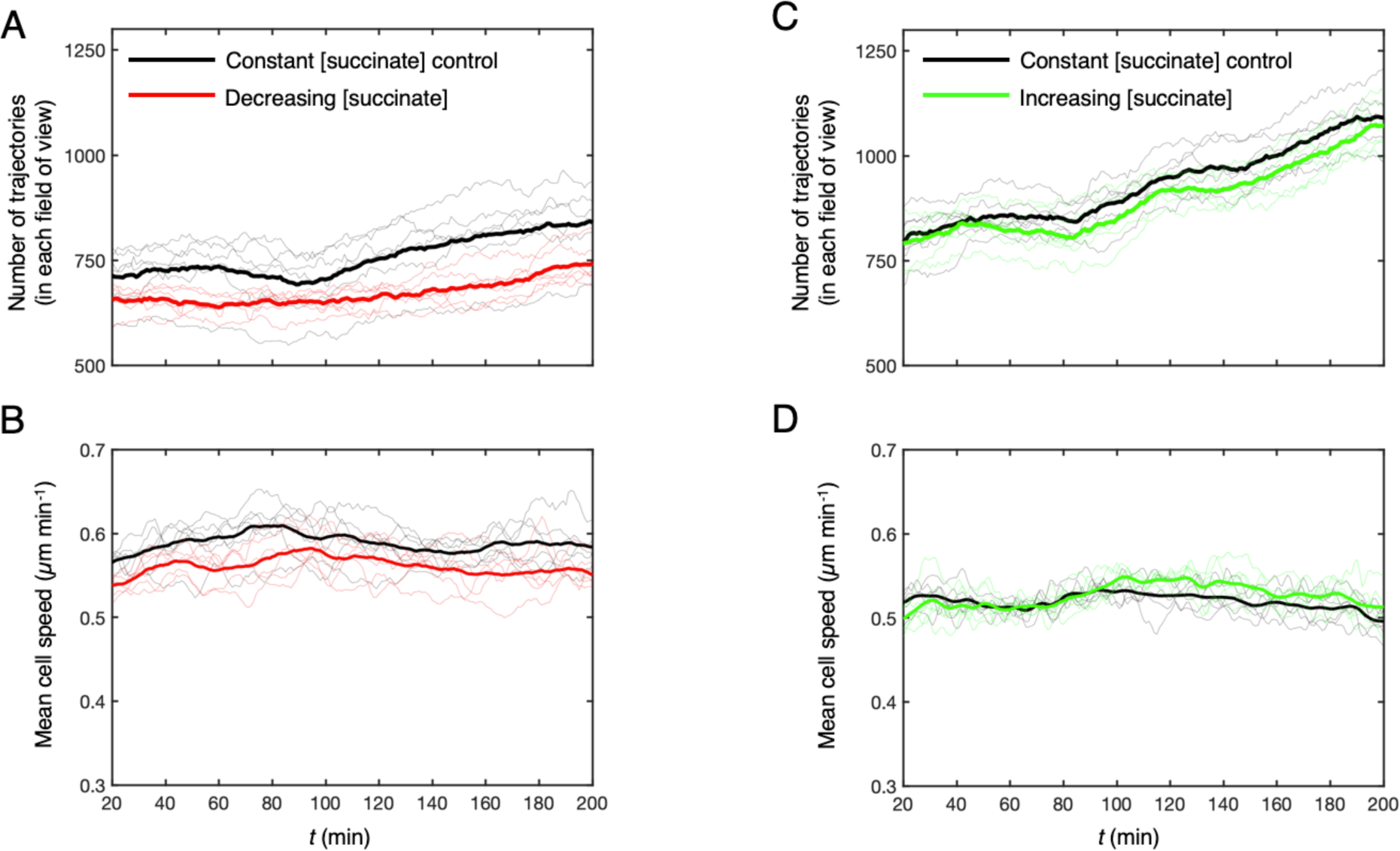
The effect of succinate on cells in our Taylor-Aris dispersion experiments. **(A)** Thin lines show the number of cell trajectories that were imaged in each of the six simultaneously imaged fields of view that were used in our Taylor-Aris dispersion experiments, whilst the thick lines show the mean. We observed that the number of cells increased gradually over the course of our approximately 3 h long experiments, regardless of whether cells were exposed to a decrease in succinate concentration over time (red lines) or to a constant concentration of succinate *C* = 1 mM in control experiments (black lines). **(B)** Cell speed remained approximately constant both in controls (black lines) and in cells exposed to decreasing succinate concentration (red lines). **(C,D)** Similar trends were observed for cells exposed to an increase (green lines) in succinate concentration over time when compared to their respective controls (black lines). The data shown here is representative of both bio-replicates.

**Figure S3.**
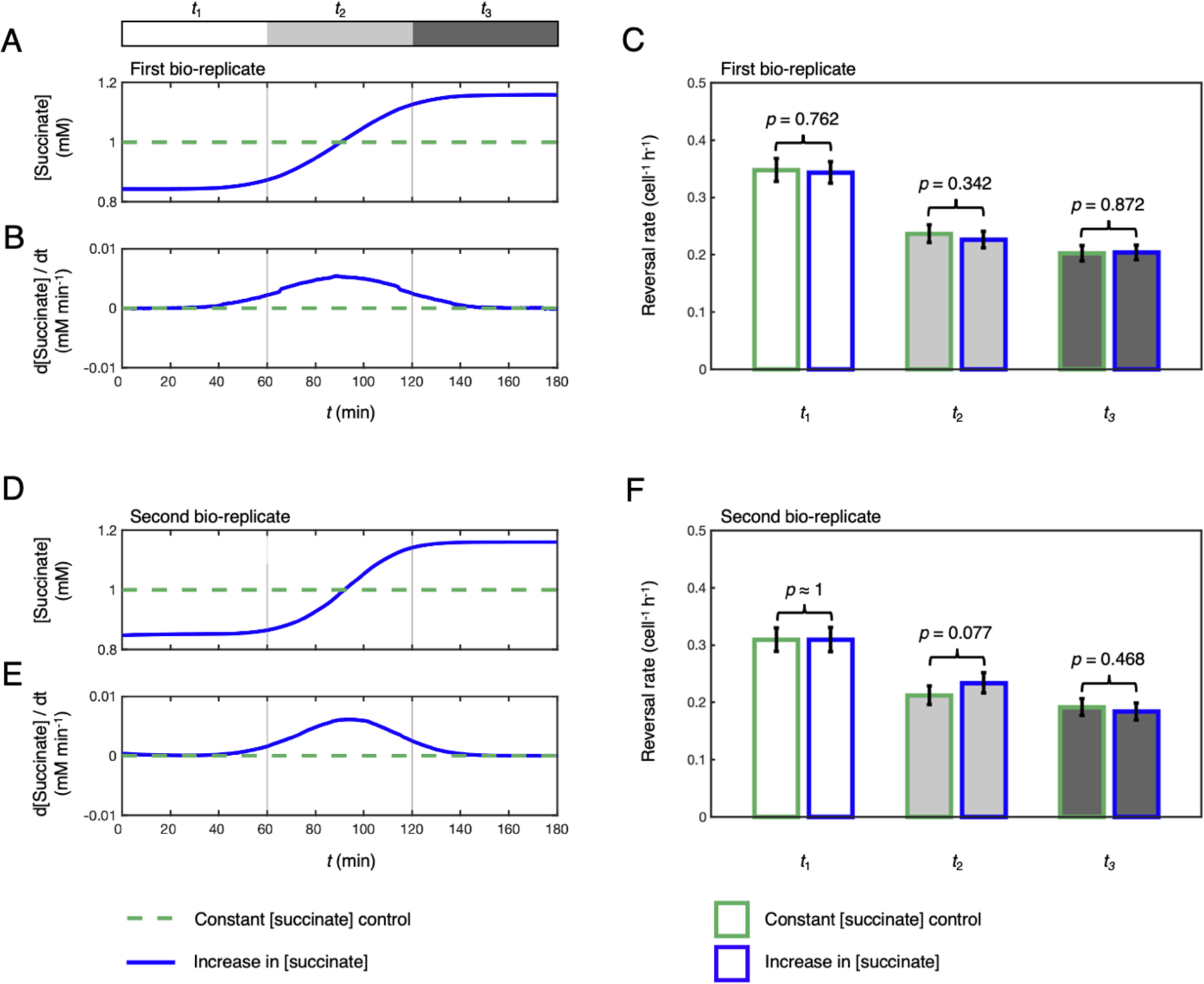
A temporal increase in succinate concentration does not induce a chemotactic response in surface-attached *P. aeruginosa*. **(A, B)** Using the same experimental approach outlined in **Fig. 2**, cells were also exposed to a temporal increase in succinate concentration from *C*MIN = 0.84 mM to *C*MAX = 1.16 mM (blue line) or to control conditions with a constant concentration *C* = 1 mM (dashed green line). This generates a mean temporal concentration gradient similar in magnitude to the gradient experienced by cells moving towards increasing succinate concentrations in the dual-inlet experiments where chemotaxis is readily observed (**Fig. S1**), but with a 16,000-fold smaller spatial gradient. If cells were able to sense these temporal stimuli, one might predict that the increase in succinate concentration over time would cause cells to suppress reversals. **(C)** Using automated reversal detection, we first confirmed that the reversal rate in the 1 h period before the succinate gradient entered the microfluidic device (time interval *t*1; white bar, blue outline), was statistically indistinguishable from the reversal rate during the same time period in a simultaneous control experiment where a constant concentration of succinate was maintained throughout (white bar, green outline). Specifically, a one-sided exact Poisson test (**Methods**) did not reject the null hypothesis that these two reversal rate measurements come from the same Poisson distribution, *p* = 0.762. Similarly, the reversal rates in the presence of a temporal succinate gradient (time interval *t*2; light grey bar, blue outline) and in the 1 h period after the gradient had cleared the microfluidic device (time interval *t*3; dark grey bar, blue outline) were statistically indistinguishable from the reversal rates during the same time periods in the control (*p* = 0.342 and *p* = 0.872). The total number of reversals observed across the six simultaneously imaged fields of view was *n*r = 2709 and 2980 across a total of *n*t = 636,364 and 709,607 trajectory points in the control and experimental conditions respectively. **(D**, **E, F)** A second bio-replicate of the experiment confirmed that when comparing between the experimental and control conditions, the reversal rates were indistinguishable during time periods *t*1 (white bars, *p* ≈ 1), *t*2 (light gray bars, *p* = 0.077) and *t3* (dark gray bars, *p* = 0.468). *n*r = 2101 and 2034 across a total of *n*t = 536,892 and 504,264 trajectory points in the control and experimental conditions respectively. Error bars show 95% confidence intervals assuming that reversals follow a Poisson distribution (**Methods**).

**Figure S4.**
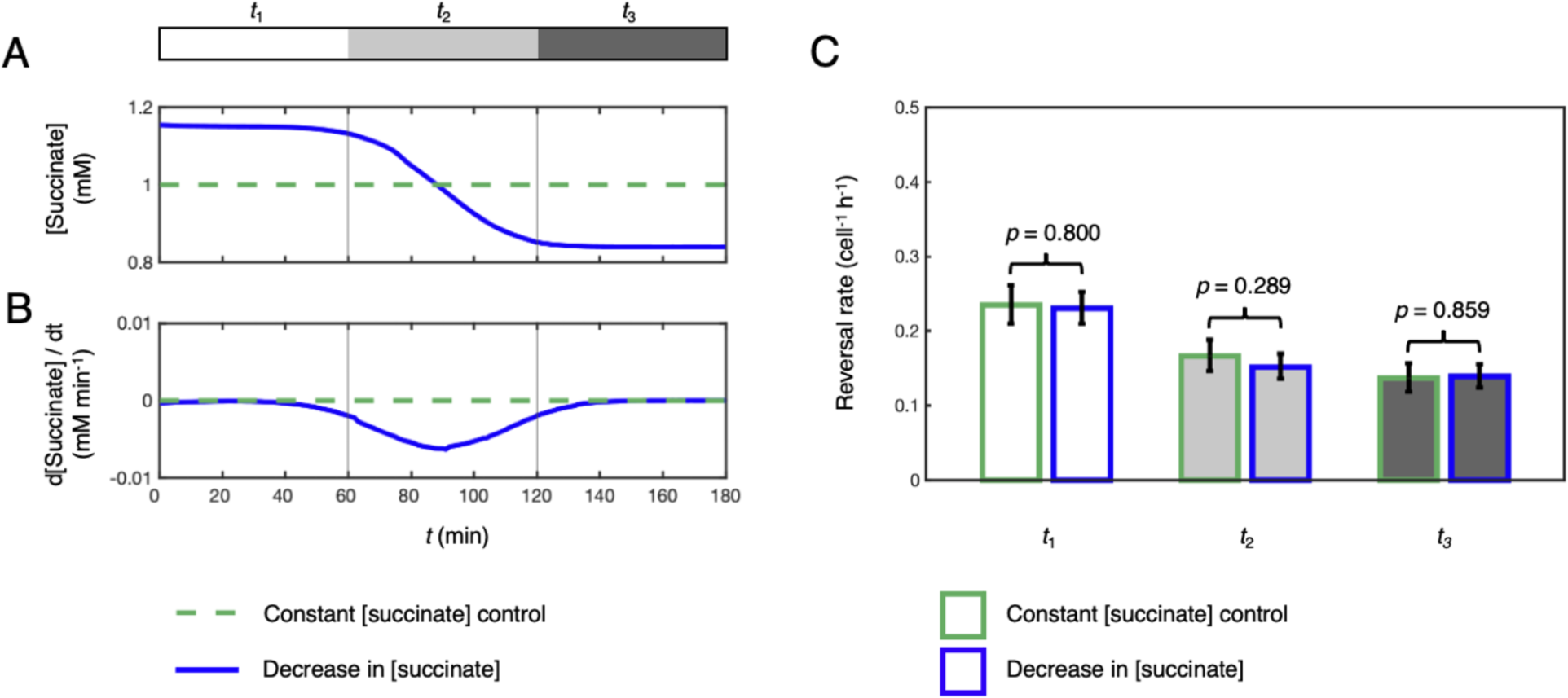
A temporal decrease in succinate concentration does not induce a chemotactic response in surface-attached *P. aeruginosa*. **(A, B)** Data shown come from a biological repeat of the experiment outlined in **Fig. 2**, where cells were either exposed to a temporal decrease in succinate concentration over time (blue lines) or to a control with a constant succinate concentration *C* = 1 mM (dashed green lines). **(C)** Using automated reversal detection, we first confirmed that the reversal rate in the 1 h period before the succinate gradient entered the microfluidic device (time interval *t*1; white bar, blue outline), was statistically indistinguishable from the reversal rate during the same time period in a simultaneous control experiment where a constant concentration of succinate was maintained throughout (white bar, green outline). Specifically, a one-sided exact Poisson test (**Methods**) did not reject the null hypothesis that these two reversal rate measurements come from the same Poisson distribution, *p* = 0.800. Similarly, the reversal rates in the presence of a temporal succinate gradient (time interval *t*2; light grey bar, blue outline) and in the 1 h period after the gradient had cleared the microfluidic device (time interval *t*3; dark grey bar, blue outline) were statistically indistinguishable from the reversal rates during the same time periods in the control (*p* = 0.289 and *p* = 0.859). The total number of reversals observed in our six simultaneously imaged fields of view was *n*r = 772 and 1072 across a total of *n*t = 259,301 and 370,801 trajectory points in the control and experimental conditions respectively. Error bars show 95% confidence intervals assuming that reversals follow a Poisson distribution (**Methods**).

**Figure S5.**
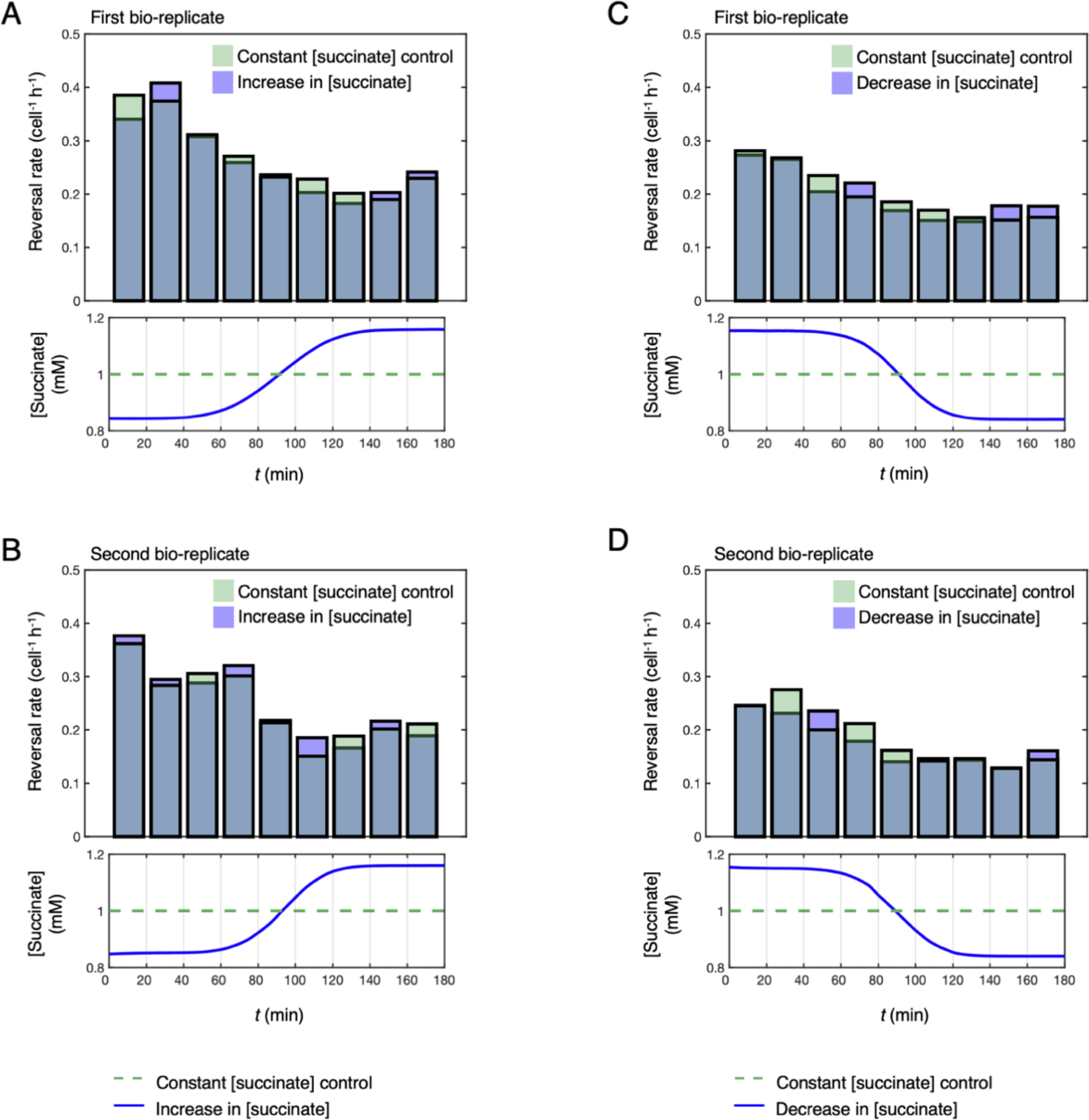
Cell reversal rate decreases over time in our Taylor-Aris dispersion experiments and in their respective controls. **(A)** In experiments that exposed cells to a temporal increase in succinate concentration (blue bars), cell reversal rate decreased over the time course of the experiments. A similar decrease was observed in the corresponding controls (green bars) where cells were exposed to a constant succinate concentration *C* = 1 mM. Similar trends were observed in a second bio-replicate of this experiment **(B)** and in two bio-replicates where cells were exposed to a temporal decrease in succinate concentration **(C, D)**.

**Figure S6.**
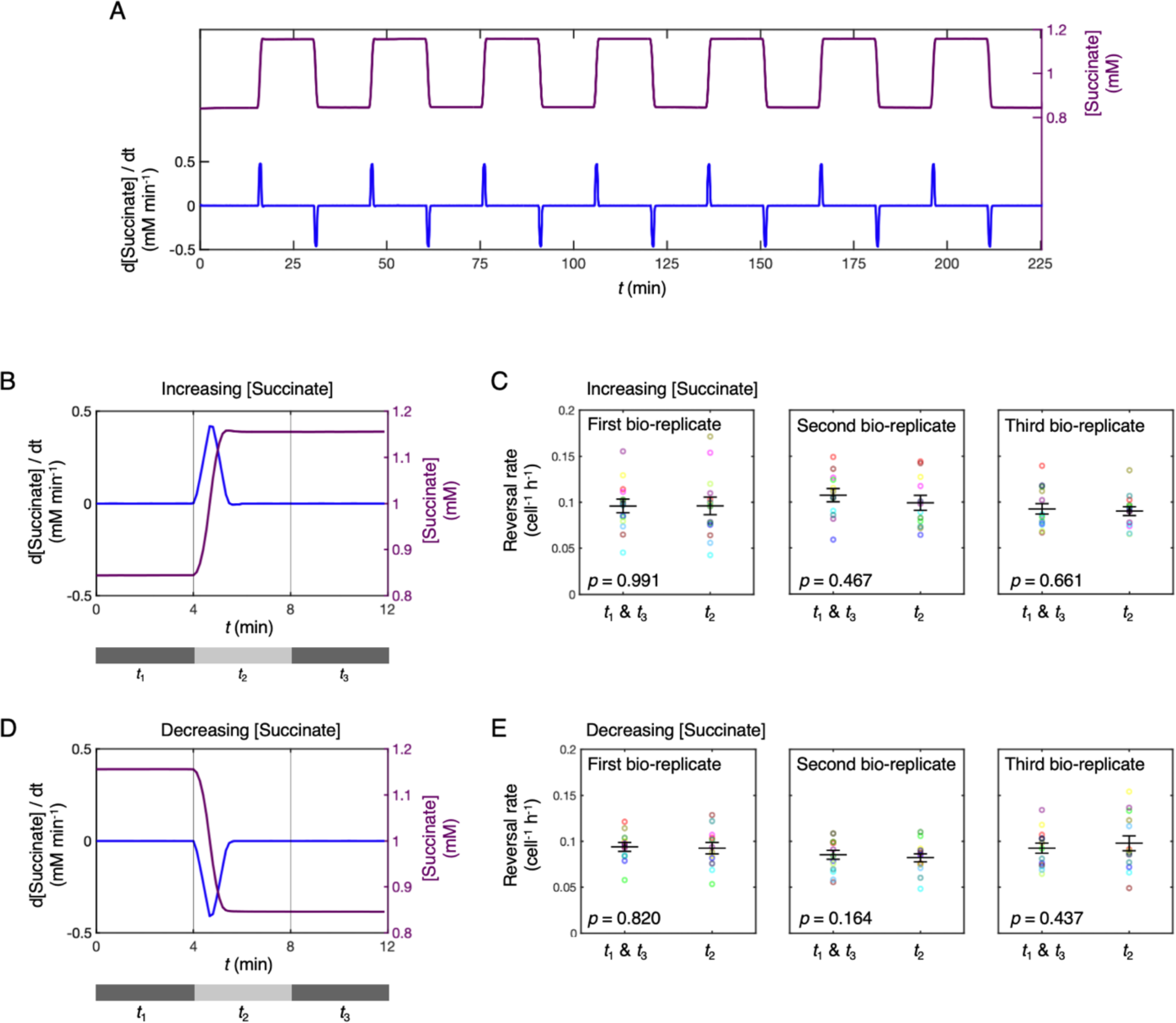
Steep and rapid temporal chemoattractant gradients do not cause surface-attached *P. aeruginosa* cells to change their reversal rate. **(A)** As discussed in the main text, twitching motility in *P. aeruginosa* is characteristically jerky and so we reasoned that cells could potentially have evolved the capacity to detect the large but ephemeral temporal changes in chemoattractant concentration caused by these intermittent displacements. To test this hypothesis, we used a microfluidic setup that exposed surface-attached cells to rapid temporal changes in succinate concentration (see **Methods**). We quantified the temporal changes in succinate concentration (purple line) and the corresponding temporal succinate gradients (blue line) that cells experienced in these experiments by labeling one of the two chemoattractant solutions with dye. In this experiment, cells are repeatedly exposed to both increases and decreases in succinate concentration. **(B)** To analyse cells’ response to these different stimuli, we first split reversal data around each increase in succinate concentration into three time-bins *t*1, *t*2, and *t*3 corresponding to the 4 min intervals before, during and after the temporal gradient. **(C)** Reversal rates were pooled across time windows *t*1 and *t*3, corresponding to time periods without any succinate concentration gradients, and compared to the reversal rates during the temporal increases in succinate concentration, time window *t*2. The mean reversal rate measured during the temporal increase in succinate concentration (large black “-” marker) was statistically indistinguishable from that when the succinate concentration was constant (a two-tailed, paired t-test of the null hypothesis of no difference in reversal rates yielded *p* = 0.991, 0.467 and 0.661 for three independent bio-replicates). Mean reversal rates were averaged across six subsequent increases in succinate concentration (see (A)) each imaged across two independent fields of view (the 12 circular markers are colour-coded to show pairs of data recorded in each of the 12 fields of view, see **Methods**). **(D, E)** Similar results were obtained when comparing reversal rates between the presence (*t*2) and absence (*t*1 and *t*3) of temporal decreases in succinate concentration (*p* = 0.820, 0.164 and 0.437 for three independent bio-replicates). Error bars show mean reversal rates plus and minus standard error.

**Figure S7.**
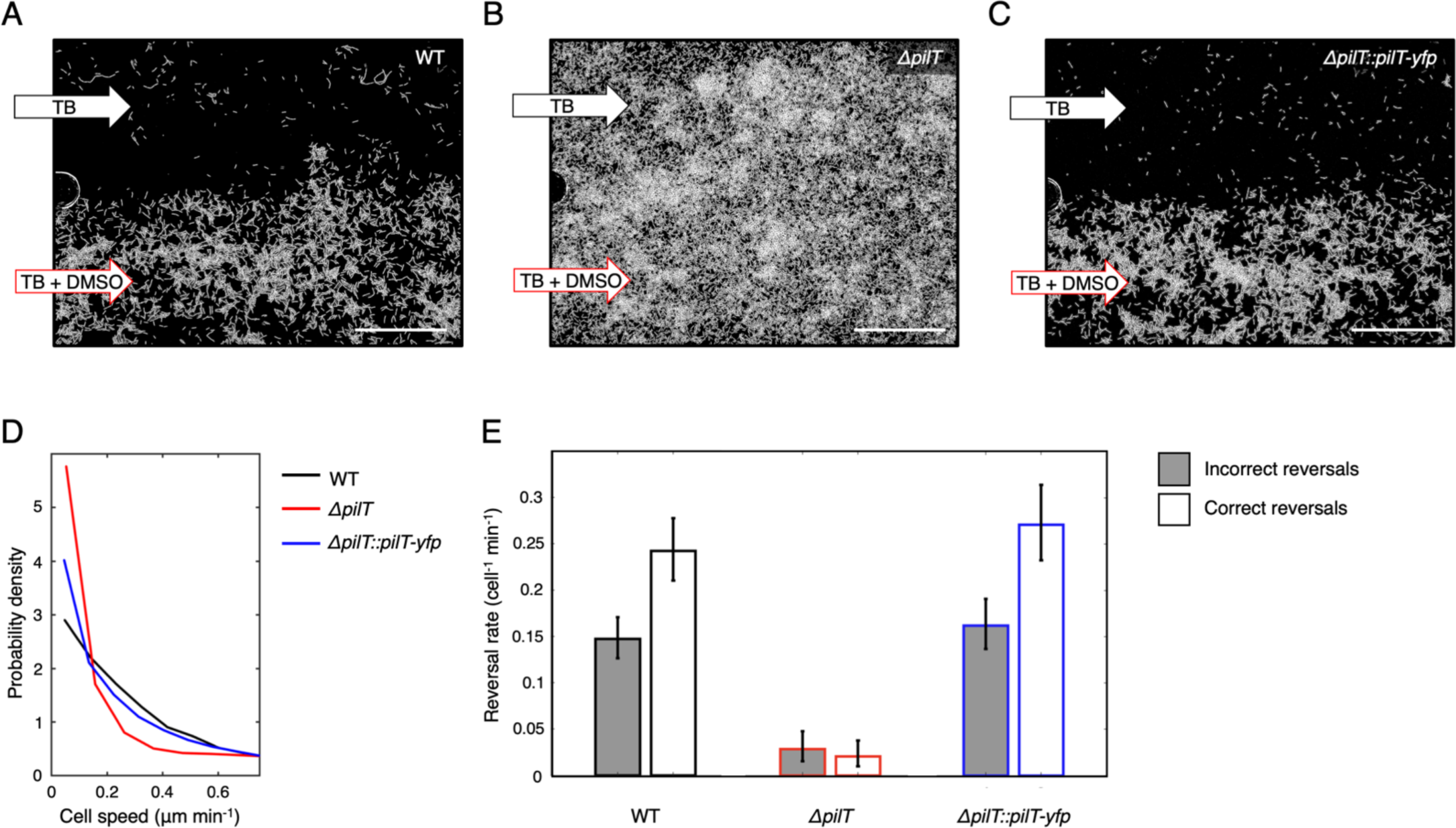
Our PilT-YFP fusion protein complements motility and chemotaxis phenotypes of a *ΔpilT* mutant. **(A)** *P. aeruginosa* cells attached to the surface of a dual-inlet microfluidic device are exposed to a concentration gradient of DMSO (a known chemoattractant [32]) by flowing TB media through one device inlet and TB media supplemented with DMSO (*C*MAX = 350 mM) through the other. WT cells (shown in white) undergo chemotaxis and accumulate in regions of high chemoattractant concentration (*t* =10 h). **(B)** *P. aeruginosa* cells lacking *pilT* (*ΔpilT*, a gene encoding a pili retraction motor) have impaired twitching motility [51] and do not undergo chemotaxis, so they distribute equally in all regions of the device (*t* = 10 h). **(C)** Our PilT-YFP translational fusion construct restores motility and chemotaxis when expressed in the *ΔpilT* strain (*ΔpilT*: PilT-YFP, *t* = 10 h). Scale bars = 50 μm. **(D)** A probability density function of cell speed (for the first 400 min of an experiment, when cells tend to exhibit their highest levels of motility) confirms that the *ΔpilT* strain (red line) has impaired twitching motility, which is restored by our PilT-YFP translational fusion (blue line). The movement speed of WT cells (black line) is shown for reference. **(E)** As discussed in the main text, WT *P. aeruginosa* cells chemotax by deploying “correct” reversals (white bars) more frequently than “incorrect” reversals (grey bars, see also **Fig. S1**). Assuming that reversals follow a Poisson distribution (**Methods**), a one-sided exact Poisson test (**Methods**) rejects the null hypothesis that the measured correct and incorrect reversal rates come from the same Poisson distribution (*p* < 0.001, total number of reversals, *n* = 382). The *ΔpilT* strain exhibits a greatly reduced reversal rate that is consistent with its impaired motility and the rate of correct reversals is indistinguishable from the rate of incorrect reversals (*p* = 0.541, *n* = 24). The WT phenotype is restored by the PilT-YFP translational fusion (*ΔpilT*: PilT-YFP; *p* < 0.001, *n* = 322). Error bars show 95% confidence intervals assuming that reversals follow a Poisson distribution (**Methods**).

**Figure S8.**
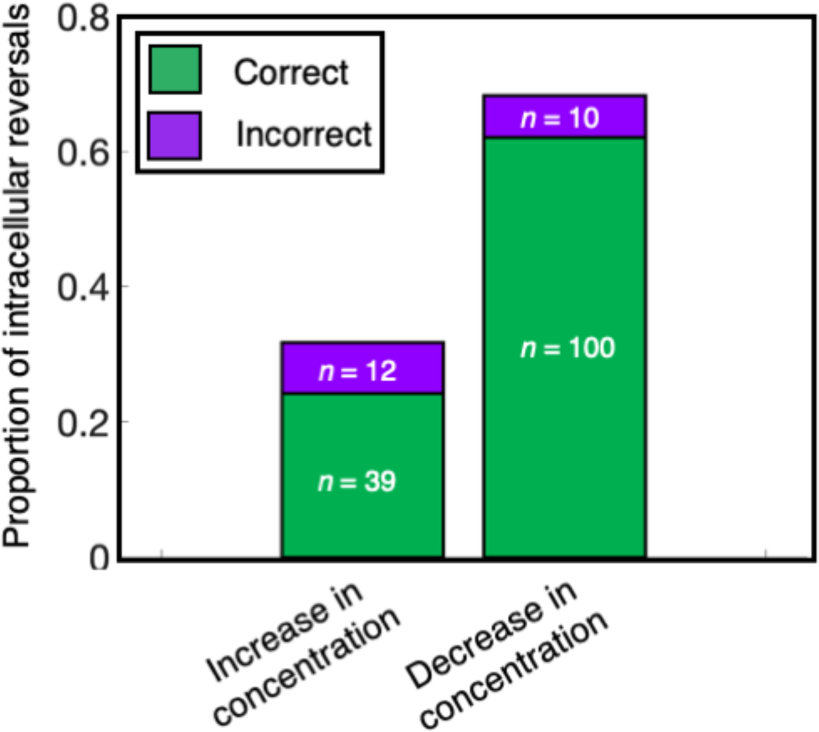
Stationary cells are more likely to undergo intracellular reversals when they have recently experienced a rapid decrease in chemoattractant concentration. In the alternating gradient experiments (**Fig. 4** and **5**), cells are exposed to large temporal changes in succinate concentration. We observed significantly more intracellular reversals in cells that experienced a rapid decrease in succinate concentration than a rapid increase. Specifically, we rejected the null hypothesis that the number of intracellular reversals following a decrease in succinate concentration is equal to the number of those following an increase in succinate concentration, (*p* = 3.83 x 10^−6^, two-tailed hypothesis test, assuming a binomial distribution with *n* = 161 trials with probability of 0.5 in each trial). This suggests that the chemotactic response depends in part on the absolute chemoattractant concentration experienced by cells [53, 54]. In both cases, “correct” intracellular reversals (green bars) were significantly more abundant than “incorrect” ones (magenta bars). Specifically, we rejected the null hypothesis that incorrect and correct intracellular reversals occurred with equal frequency when the concentration was increasing (*p* = 1.98 x 10^−4^, one-tailed hypothesis test assuming a binomial process with *n* = 51 trials and probability of 0.5 in each trial) and for when the concentration was decreasing (*p* = 8.01 x 10^−20^, one-tailed hypothesis test assuming a binomial distribution with *n* = 110 trials and probability of 0.5 in each trial). Lastly, we note that cells experiencing a decrease in succinate concentration were significantly more likely to perform correct reversals than those experiencing an increase in succinate concentration. This analysis used a two-tailed Fisher’s exact test to reject the null hypothesis that there was no association between the sign of the temporal succinate gradient and whether the intracellular reversal was correct or incorrect (*p* = 0.024).

## Bibliography

1. Jin T, Xu X, Hereld D: Chemotaxis, chemokine receptors and human disease. Cytokine 2008, 44(1):1–8.

2. Matilla MA, Krell T: The effect of bacterial chemotaxis on host infection and pathogenicity. FEMS Microbiol Rev 2018, 42(1):40–67.

3. Stocker R, Seymour JR: Ecology and physics of bacterial chemotaxis in the ocean. Microbiol Mol Biol Rev 2012, 76(4):792–812.

4. Stocker R, Seymour JR, Samadani A, Hunt DE, Polz MF: Rapid chemotactic response enables marine bacteria to exploit ephemeral microscale nutrient patches. Proc Natl Acad Sci U S A 2008, 105(11):4209–4214.

5. Friedrich BM, Julicher F: Chemotaxis of sperm cells. Proc Natl Acad Sci U S A 2007, 104(33):13256–13261.

6. Nichols JM, Veltman D, Kay RR: Chemotaxis of a model organism: progress with Dictyostelium. Curr Opin Cell Biol 2015, 36:7–12.

7. Dormann D, Weijer CJ: Chemotactic cell movement during development. Curr Opin Genet Dev 2003, 13(4):358–364.

8. Iglesias PA, Devreotes PN: Navigating through models of chemotaxis. Curr Opin Cell Biol 2008, 20(1):35–40.

9. Berg HC: E. coli in motion: Springer, New York; 2004.

10. Bi S, Sourjik V: Stimulus sensing and signal processing in bacterial chemotaxis. Curr Opin Microbiol 2018, 45:22–29.

11. Wadhams GH, Armitage JP: Making sense of it all: bacterial chemotaxis. Nat Rev Mol Cell Biol 2004, 5(12):1024–1037.

12. Hellingwerf KJ, Postma PW, Tommassen J, Westerhoff HV: Signal transduction in bacteria: phospho-neural network(s) in Escherichia coli? FEMS Microbiol Rev 1995, 16(4):309–321.

13. Zschiedrich CP, Keidel V, Szurmant H: Molecular mechanisms of two-component signal transduction. J Mol Biol 2016, 428(19):3752–3775.

14. Baker MD, Wolanin PM, Stock JB: Signal transduction in bacterial chemotaxis. Bioessays 2006, 28(1):9–22.

15. Porter SL, Wadhams GH, Armitage JP: Rhodobacter sphaeroides: complexity in chemotactic signalling. Trends Microbiol 2008, 16(6):251–260.

16. Bischoff DS, Ordal GW: Bacillus subtilis chemotaxis: a deviation from the Escherichia coli paradigm. Mol Microbiol 1992, 6(1):23–28.

17. Thar R, Kuhl M: Bacteria are not too small for spatial sensing of chemical gradients: an experimental evidence. Proc Natl Acad Sci U S A 2003, 100(10):5748–5753.

18. Jin T: Gradient sensing during chemotaxis. Curr Opin Cell Biol 2013, 25(5):532–537.

19. Chen YE, Tropini C, Jonas K, Tsokos CG, Huang KC, Laub MT: Spatial gradient of protein phosphorylation underlies replicative asymmetry in a bacterium. Proc Natl Acad Sci U S A 2011, 108(3):1052–1057.

20. Berg HC, Purcell EM: Physics of chemoreception. Biophys J 1977, 20(2):193–219.

21. Tindall MJ, Gaffney EA, Maini PK, Armitage JP: Theoretical insights into bacterial chemotaxis. Wiley Interdiscip Rev Syst Biol Med 2012, 4(3):247–259.

22. Stephens BB, Loar SN, Alexandre G: Role of CheB and CheR in the complex chemotactic and aerotactic pathway of Azospirillum brasilense. J Bacteriol 2006, 188(13):4759–4768.

23. Kato J, Kim HE, Takiguchi N, Kuroda A, Ohtake H: Pseudomonas aeruginosa as a model microorganism for investigation of chemotactic behaviors in ecosystem. J Biosci Bioeng 2008, 106(1):1–7.

24. Packer HL, Gauden DE, Armitage JP: The behavioural response of anaerobic Rhodobacter sphaeroides to temporal stimuli. Microbiology (Reading) 1996, 142 (Pt 3):593–599.

25. Macnab RM, Koshland DE, Jr.: The gradient-sensing mechanism in bacterial chemotaxis. Proc Natl Acad Sci U S A 1972, 69(9):2509–2512.

26. Rao CV, Glekas GD, Ordal GW: The three adaptation systems of Bacillus subtilis chemotaxis. Trends Microbiol 2008, 16(10):480–487.

27. Brown DA, Berg HC: Temporal stimulation of chemotaxis in Escherichia coli. Proc Natl Acad Sci U S A 1974, 71(4):1388–1392.

28. Lefevre CT, Bennet M, Landau L, Vach P, Pignol D, Bazylinski DA, Frankel RB, Klumpp S, Faivre D: Diversity of magneto-aerotactic behaviors and oxygen sensing mechanisms in cultured magnetotactic bacteria. Biophys J 2014, 107(2):527–538.

29. Hall-Stoodley L, Costerton JW, Stoodley P: Bacterial biofilms: from the natural environment to infectious diseases. Nat Rev Microbiol 2004, 2(2):95–108.

30. Costerton JW, Lewandowski Z, Caldwell DE, Korber DR, Lappin-Scott HM: Microbial biofilms. Annu Rev Microbiol 1995, 49:711–745.

31. Flemming HC, Wuertz S: Bacteria and archaea on Earth and their abundance in biofilms. Nat Rev Microbiol 2019, 17(4):247–260.

32. Oliveira NM, Foster KR, Durham WM: Single-cell twitching chemotaxis in developing biofilms. Proc Natl Acad Sci U S A 2016, 113(23):6532–6537.

33. Conrad JC, Gibiansky ML, Jin F, Gordon VD, Motto DA, Mathewson MA, Stopka WG, Zelasko DC, Shrout JD, Wong GC: Flagella and pili-mediated near-surface single-cell motility mechanisms in P. aeruginosa. Biophys J 2011, 100(7):1608–1616.

34. Meacock OJ, Doostmohammadi A, Foster KR, Yeomans JM, Durham WM: Bacteria solve the problem of crowding by moving slowly. Nature Physics 2021, 17(3):205–210.

35. Harshey RM: Bacterial motility on a surface: many ways to a common goal. Annu Rev Microbiol 2003, 57:249–273.

36. Jarrell KF, McBride MJ: The surprisingly diverse ways that prokaryotes move. Nat Rev Microbiol 2008, 6(6):466–476.

37. Burrows LL: Pseudomonas aeruginosa twitching motility: type IV pili in action. Annu Rev Microbiol 2012, 66:493–520.

38. Cai Q, Li Z, Ouyang Q, Luo C, Gordon VD: Singly flagellated Pseudomonas aeruginosa chemotaxes efficiently by unbiased motor regulation. mBio 2016, 7(2):e00013.

39. Maier B, Wong GCL: How bacteria use type IV pili machinery on surfaces. Trends Microbiol 2015, 23(12):775–788.

40. Kuhn MJ, Tala L, Inclan YF, Patino R, Pierrat X, Vos I, Al-Mayyah Z, Macmillan H, Negrete J, Jr., Engel JN et al: Mechanotaxis directs Pseudomonas aeruginosa twitching motility. Proc Natl Acad Sci U S A 2021, 118(30):e2101759118.

41. Jin F, Conrad JC, Gibiansky ML, Wong GC: Bacteria use type-IV pili to slingshot on surfaces. Proc Natl Acad Sci U S A 2011, 108(31):12617–12622.

42. Schuergers N, Nurnberg DJ, Wallner T, Mullineaux CW, Wilde A: PilB localization correlates with the direction of twitching motility in the cyanobacterium Synechocystis sp. PCC 6803. Microbiology (Reading) 2015, 161(Pt 5):960–966.

43. Taylor G: Dispersion of soluble matter in solvent flowing slowly through a tube. Proc R Soc Lon Ser-A 1953, 219(1137):186–203.

44. Bello MS, Rezzonico R, Righetti PG: Use of taylor-aris dispersion for measurement of a solute diffusion coefficient in thin capillaries. Science 1994, 266(5186):773-776.

45. O’Toole GA, Wong GC: Sensational biofilms: surface sensing in bacteria. Curr Opin Microbiol 2016, 30:139–146.

46. Zhao K, Tseng BS, Beckerman B, Jin F, Gibiansky ML, Harrison JJ, Luijten E, Parsek MR, Wong GCL: Psl trails guide exploration and microcolony formation in Pseudomonas aeruginosa biofilms. Nature 2013, 497(7449):388-391.

47. Schumacher D, Sogaard-Andersen L: Regulation of cell polarity in motility and cell division in Myxococcus xanthus. Annu Rev Microbiol 2017, 71:61–78.

48. Bulyha I, Schmidt C, Lenz P, Jakovljevic V, Hone A, Maier B, Hoppert M, Sogaard-Andersen L: Regulation of the type IV pili molecular machine by dynamic localization of two motor proteins. Mol Microbiol 2009, 74(3):691–706.

49. Leonardy S, Miertzschke M, Bulyha I, Sperling E, Wittinghofer A, Sogaard-Andersen L: Regulation of dynamic polarity switching in bacteria by a Ras-like G-protein and its cognate GAP. EMBO J 2010, 29(14):2276–2289.

50. Wu Y, Kaiser AD, Jiang Y, Alber MS: Periodic reversal of direction allows Myxobacteria to swarm. Proc Natl Acad Sci U S A 2009, 106(4):1222–1227.

51. Bertrand JJ, West JT, Engel JN: Genetic analysis of the regulation of type IV pilus function by the Chp chemosensory system of Pseudomonas aeruginosa. J Bacteriol 2010, 192(4):994–1010.

52. Koch MD, Fei C, Wingreen NS, Shaevitz JW, Gitai Z: Competitive binding of independent extension and retraction motors explains the quantitative dynamics of type IV pili. Proc Natl Acad Sci U S A 2021, 118(8).

53. Adler M, Alon U: Fold-change detection in biological systems. Curr Op Sys Biol 2018, 8:81–89.

54. Rivero MA, Tranquillo RT, Buettner HM, Lauffenburger DA: Transport models for chemotactic cell-populations based on individual cell behavior. Chem Eng Sci 1989, 44(12):2881–2897.

55. Rickert P, Weiner OD, Wang F, Bourne HR, Servant G: Leukocytes navigate by compass: roles of PI3Kγ and its lipid products. Trends Cell Biol 2000, 10(11):466–473.

56. Swaney KF, Huang CH, Devreotes PN: Eukaryotic chemotaxis: a network of signaling pathways controls motility, directional sensing, and polarity. Annu Rev Biophys 2010, 39:265–289.

57. Mika JT, Poolman B: Macromolecule diffusion and confinement in prokaryotic cells. Curr Opin Biotechnol 2011, 22(1):117–126.

58. Mullineaux CW, Nenninger A, Ray N, Robinson C: Diffusion of green fluorescent protein in three cell environments in Escherichia coli. J Bacteriol 2006, 188(10):3442–3448.

59. Dusenbery DB: Spatial sensing of stimulus gradients can be superior to temporal sensing for free-swimming bacteria. Biophys J 1998, 74(5):2272–2277.

60. Zobel S, Benedetti I, Eisenbach L, de Lorenzo V, Wierckx N, Blank LM: Tn7-based device for calibrated heterologous gene expression in Pseudomonas putida. ACS Synth Biol 2015, 4(12):1341–1351.

61. Salis HM, Mirsky EA, Voigt CA: Automated design of synthetic ribosome binding sites to control protein expression. Nat Biotechnol 2009, 27(10):946–950.

62. Chen X, Zaro JL, Shen WC: Fusion protein linkers: property, design and functionality. Adv Drug Deliv Rev 2013, 65(10):1357–1369.

63. Choi KH, Schweizer HP: mini-Tn7 insertion in bacteria with single attTn7 sites: example Pseudomonas aeruginosa. Nat Protoc 2006, 1(1):153–161.

64. Haubert K, Drier T, Beebe D: PDMS bonding by means of a portable, low-cost corona system. Lab Chip 2006, 6(12):1548–1549.

65. Schindelin J, Arganda-Carreras I, Frise E, Kaynig V, Longair M, Pietzsch T, Preibisch S, Rueden C, Saalfeld S, Schmid B et al: Fiji: an open-source platform for biological-image analysis. Nat Methods 2012, 9(7):676–682.

66. Miura K: Bleach correction ImageJ plugin for compensating the photobleaching of time-lapse sequences. F1000Res 2020, 9:1494.

67. Tinevez JY, Perry N, Schindelin J, Hoopes GM, Reynolds GD, Laplantine E, Bednarek SY, Shorte SL, Eliceiri KW: TrackMate: An open and extensible platform for single-particle tracking. Methods 2017, 115:80–90.

68. Meacock OJ, Durham WM: Tracking bacteria at high density with FAST, the Feature-Assisted Segmenter/Tracker. bioRxiv 2023:2021.2011.2026.470050.

69. Carter T, Buensuceso RN, Tammam S, Lamers RP, Harvey H, Howell PL, Burrows LL: The type IVa pilus machinery is recruited to sites of future cell division. mBio 2017, 8(1):e02103–02116.

70. DeLisi C, Marchetti F, Del Grosso G: A theory of measurement error and its implications for spatial and temporal gradient sensing during chemotaxis. Cell Biophys 1982, 4(2-3):211–229.

