## Supplementary File 1 for "Bacteria use spatial sensing to direct chemotaxis on surfaces"

| | CELL IDENTIFIERS | | 0 = incorrect intracellular reversal<br>1 = correct intracellular reversal | 0 = nonpolar<br>1 = unipolar<br>2 = bipolar<br>na = not assignable | 0 = cell experiences increasing [succinate]<br>1= cell experiences decreasing [succinate]<br>na = not assignable |
|---|---|---|---|---|---|
| **MOVIE** | **BIOREP** | **INTRACELLULAR REVERSAL** | **CORRECT OR INCORRECT** | **INITIAL POLARITY** | **TEMPORAL CHANGE IN [SUCCINATE]** |
| 1 | 1 | 1 | 1 | 0 | 1 |
| | | 2 | 1 | 2 | 0 |
| | | 3 | 0 | 0 | 1 |
| 2 | 1 | 1 | 1 | 0 | 1 |
| | | 2 | 1 | 0 | 1 |
| | | 3 | 1 | 0 | 0 |
| | | 4 | 1 | 0 | 0 |
| | | 5 | 1 | 0 | 1 |
| | | 6 | 1 | 1 | 1 |
| | | 7 | 1 | 1 | 0 |
| | | 8 | 1 | 2 | 0 |
| | | 9 | 1 | 2 | 0 |
| | | 10 | 1 | 0 | 0 |
| 3 | 1 | 1 | 1 | 2 | na |
| | | 2 | 1 | 1 | 1 |
| | | 3 | 1 | 0 | 1 |
| | | 4 | 1 | 0 | 0 |
| | | 5 | 1 | 1 | 0 |
| | | 6 | 1 | 0 | 0 |
| | | 7 | 1 | 0 | 0 |
| | | 8 | 1 | 1 | 0 |
| 4 | 1 | 1 | 0 | 0 | 0 |
| | | 2 | 1 | 1 | 1 |
| | | 3 | 0 | 0 | 0 |
| | | 4 | 1 | 1 | 0 |
| | | 5 | 1 | 1 | 0 |
| | | 6 | 1 | 2 | na |
| | | 7 | 1 | 0 | 0 |
| 5 | 1 | 1 | 1 | 1 | 0 |
| | | 2 | 1 | 0 | 0 |
| | | 3 | 1 | 2 | 0 |
| | | 4 | 1 | 2 | 0 |
| | | 5 | 1 | 2 | 0 |
| | | 6 | 1 | 2 | 1 |
| | | 7 | 1 | 1 | 0 |
| | | 8 | 0 | 2 | 1 |
| | | 9 | 1 | 2 | 0 |
| | | 10 | 1 | 1 | 1 |
| | | 11 | 1 | 2 | 0 |
| | | 12 | 1 | 1 | 0 |
| | | 13 | 1 | 2 | 0 |
| | | 14 | 1 | 0 | 1 |
| 6 | 1 | 1 | 0 | 2 | 1 |
| | | 2 | 1 | 1 | 1 |
| | | 3 | 1 | 2 | 0 |
| | | 4 | 1 | 0 | 0 |
| | | 5 | 1 | 0 | 0 |
| | | 6 | 1 | 0 | 0 |
| | | 7 | 0 | 0 | 1 |
| | | 8 | 1 | 1 | 1 |

| MOVIE | BIOREP | INTRACELLULAR REVERSAL | CORRECT OR INCORRECT | INITIAL POLARITY | TEMPORAL CHANGE IN [SUCCINATE] |
|---|---|---|---|---|---|
| | | 9 | 1 | 0 | 1 |
| | | 10 | 1 | 2 | 0 |
| | | 11 | 1 | 2 | 0 |
| | | 12 | 1 | 0 | 0 |
| | | 13 | 1 | 2 | 1 |
| 7 | 1 | 1 | 1 | na | 1 |
| | | 2 | 1 | 0 | 1 |
| | | 3 | 1 | 2 | 0 |
| | | 4 | 1 | 1 | 0 |
| | | 5 | 1 | 0 | 0 |
| | | 6 | 1 | 1 | 0 |
| | | 7 | 1 | 1 | 0 |
| 8 | 1 | 1 | 1 | 1 | 0 |
| | | 2 | 1 | 0 | 1 |
| | | 3 | 1 | 1 | 0 |
| | | 4 | 1 | na | 0 |
| | | 5 | 1 | 1 | 0 |
| | | 6 | 1 | 0 | 0 |
| | | 7 | 1 | 2 | 0 |
| 9 | 1 | 1 | 0 | 2 | 1 |
| | | 2 | 1 | 0 | 1 |
| | | 3 | 1 | 0 | 1 |
| | | 4 | 1 | 1 | 0 |
| | | 5 | 1 | 1 | na |
| | | 6 | 0 | 2 | 0 |
| | | 7 | 1 | 1 | 0 |
| | | 8 | 0 | 1 | 0 |
| | | 9 | 1 | 2 | 0 |
| | | 10 | 1 | 1 | 1 |
| | | 11 | 1 | 2 | 1 |
| | | 12 | 1 | 0 | 1 |
| | | 13 | 1 | 1 | na |
| | | 14 | 1 | 1 | 0 |
| | | 15 | 1 | 1 | 0 |
| | | 16 | 1 | 1 | 1 |
| | | 17 | 0 | 2 | 1 |
| | | 18 | 0 | 2 | 0 |
| | | 19 | 1 | 2 | 0 |
| | | 20 | 1 | 1 | na |
| | | 21 | 1 | 2 | 0 |
| | | 22 | 1 | 0 | 1 |
| | | 23 | 1 | 1 | na |
| | | 24 | 1 | 0 | 0 |
| | | 25 | 1 | 2 | 0 |
| | | 26 | 1 | 0 | 1 |
| 10 | 2 | 1 | 1 | 0 | 0 |
| | | 2 | 1 | 2 | 0 |
| | | 3 | 0 | 1 | 1 |
| | | 4 | 1 | 0 | 1 |
| | | 5 | 1 | 0 | 1 |
| | | 6 | 1 | 2 | 0 |
| | | 7 | 1 | 0 | 0 |
| | | 8 | 1 | 1 | 0 |

| | A | B | C | D | E | F |
|---|---|---|---|---|---|---|
| 111 | MOVIE | BIOREP | INTRACELLULAR REVERSAL | CORRECT OR INCORRECT | INITIAL POLARITY | TEMPORAL CHANGE IN [SUCCINATE] |
| 112 | | | | | | |
| 113 | | | 9 | 1 | 0 | 0 |
| 114 | | | 10 | 1 | 0 | 0 |
| 115 | | | 11 | 1 | 1 | 0 |
| 116 | | | 12 | 1 | 1 | 0 |
| 117 | | | 13 | 1 | 2 | 0 |
| 118 | | | 14 | 1 | 2 | 1 |
| 119 | | | 15 | 1 | 2 | 0 |
| 120 | 11 | 2 | 1 | 1 | 1 | 1 |
| 121 | | | 2 | 1 | 2 | 0 |
| 122 | | | 3 | 1 | 2 | 0 |
| 123 | | | 4 | 1 | 2 | 0 |
| 124 | | | 5 | 1 | 2 | 0 |
| 125 | | | 6 | 1 | 1 | 1 |
| 126 | | | 7 | 1 | 2 | 0 |
| 127 | 12 | 3 | 1 | 0 | 2 | 1 |
| 128 | | | 2 | 1 | 1 | 1 |
| 129 | | | 3 | 1 | 0 | 0 |
| 130 | | | 4 | 1 | 2 | 0 |
| 131 | | | 5 | 1 | 2 | 1 |
| 132 | | | 6 | 1 | 2 | 1 |
| 133 | | | 7 | 1 | 2 | 0 |
| 134 | | | 8 | 1 | 2 | 0 |
| 135 | | | 9 | 1 | 2 | 0 |
| 136 | | | 10 | 1 | 2 | 0 |
| 137 | | | 11 | 0 | 0 | 1 |
| 138 | | | 12 | 1 | 0 | 0 |
| 139 | | | 13 | 1 | 1 | 0 |
| 140 | | | 14 | 1 | 2 | 0 |
| 141 | | | 15 | 1 | 2 | 0 |
| 142 | | | 16 | 1 | 2 | 0 |
| 143 | 13 | 3 | 1 | 1 | 0 | 1 |
| 144 | | | 2 | 1 | 2 | na |
| 145 | | | 3 | 1 | 2 | 0 |
| 146 | | | 4 | 1 | 2 | 0 |
| 147 | | | 5 | 0 | 0 | 1 |
| 148 | | | 6 | 1 | 0 | 1 |
| 149 | | | 7 | 1 | 1 | 0 |
| 150 | 14 | 3 | 1 | 0 | 2 | 1 |
| 151 | | | 2 | 0 | 2 | 0 |
| 152 | | | 3 | 0 | 2 | 0 |
| 153 | | | 4 | 1 | 2 | 0 |
| 154 | | | 5 | 1 | 0 | 0 |
| 155 | | | 6 | 1 | 2 | 0 |
| 156 | | | 7 | 1 | 2 | 0 |
| 157 | | | 8 | 1 | 2 | 0 |
| 158 | | | 9 | 1 | 2 | 0 |
| 159 | 15 | 3 | 1 | 0 | 1 | na |
| 160 | | | 2 | 1 | 1 | 0 |
| 161 | | | 3 | 1 | 0 | 0 |
| 162 | | | 4 | 1 | 2 | 0 |
| 163 | | | 5 | 0 | 1 | 1 |
| 164 | | | 6 | 1 | 2 | 0 |
| 165 | | | 7 | 0 | 2 | 0 |

| | A | B | C | D | E | F |
|---|---|---|---|---|---|---|
| 166 | **MOVIE** | **BIOREP** | **INTRACELLULAR REVERSAL** | **CORRECT OR INCORRECT** | **INITIAL POLARITY** | **TEMPORAL CHANGE IN [SUCCINATE]** |
| 167 | | | | | | |
| 168 | | | **8** | 1 | 2 | 0 |
| 169 | | | **9** | 1 | 1 | na |
| 170 | | | **10** | 1 | 2 | 0 |
| 171 | | | **11** | 1 | 2 | 0 |
| 172 | | | **12** | 1 | 2 | 0 |
| 173 | | | **13** | 1 | 1 | 0 |
| 174 | | | **14** | 1 | 0 | na |
| 175 | **16** | **3** | **1** | 1 | 1 | 1 |
| 176 | | | **2** | 1 | 0 | 0 |
| 177 | | | **3** | 0 | 0 | 0 |
| 178 | | | **4** | 0 | 0 | 0 |
| 179 | | | **5** | 1 | 1 | 0 |
| 180 | | | **6** | 1 | 0 | 0 |
| 181 | | | **7** | 1 | 1 | 1 |
| 182 | | | **8** | 1 | 0 | 1 |
