## Appendix Movies - Legend for "Bacteria use spatial sensing to direct chemotaxis on surfaces"

**Appendix.** Each of the intracellular reversals included in our analyses (*n* = 171) can be observed in the 16 different supplementary movies included in the appendix. In each movie, the succinate gradient is shown in blue with the arrow at the bottom indicating the direction in which the succinate concentration increases. The flow moves vertically from top to bottom. The movie pauses on the frame before the succinate gradient changes direction to mark the cells that subsequently perform an intracellular reversal. The shape of the symbol used to mark the cells corresponds to the cell's initial FimX-YFP polarity prior to the change in gradient orientation (see legend) and symbol colour corresponds to whether the intracellular reversal is "correct" (green) or "incorrect" (magenta). A summary of each intracellular reversal and how they were classified is provided in **Supplementary File 1** and the detailed set of rules that was used to detect and classify intracellular reversals are outlined in the **Methods**. For clarity, we only show the YFP fluorescent images taken at 2.5 min intervals and have omitted the brightfield images that were taken at 8 sec intervals.
